# Developmental transcriptional control of mitochondrial homeostasis is required for activity-dependent synaptic connectivity

**DOI:** 10.1101/2023.06.11.544500

**Authors:** Iryna Mohylyak, Mercedes Bengochea, Carlos Pascual-Caro, Noemi Asfogo, Sara Fonseca-Topp, Natasha Danda, Zeynep Kalender Atak, Maxime De Waegeneer, Pierre-Yves Plaçais, Thomas Preat, Stein Aerts, Olga Corti, Jaime de Juan-Sanz, Bassem A. Hassan

**Affiliations:** Institut du Cerveau-Paris Brain Institute (ICM), Sorbonne Université, Inserm, CNRS, Hôpital Pitié-Salpêtrière, 75013 Paris, France; Laboratory of Computational Biology, Center for Brain and Disease Research – VIB-KU Leuven, Leuven, 3000, Belgium; Energy & Memory Brain Plasticity (UMR 8249), CNRS, ESPCI Paris, PSL Research University, 10 rue Vauquelin, 75005 Paris, France

**Keywords:** Brain development, neuronal circuit wiring, transcription factor, dTzap/TZAP, mitochondria, synaptic release, Drosophila

## Abstract

During neuronal circuit formation, local control of axonal organelles ensures proper synaptic connectivity. Whether this process is genetically encoded is unclear and if so, its developmental regulatory mechanisms remain to be identified. We hypothesized that developmental transcription factors regulate critical parameters of organelle homeostasis that contribute to circuit wiring. We combined cell type-specific transcriptomics with a genetic screen to discover such factors. We identified Telomeric Zinc finger-Associated Protein (TZAP) as a temporal developmental regulator of neuronal mitochondrial homeostasis genes, including *Pink1*. In *Drosophila*, loss of *dTzap* function during visual circuit development leads to loss of activity-dependent synaptic connectivity, that can be rescued by *Pink1* expression. At the cellular level, loss of *dTzap/TZAP* leads to defects in mitochondrial morphology, attenuated calcium uptake and reduced synaptic vesicle release in fly and mammalian neurons. Our findings highlight developmental transcriptional regulation of mitochondrial homeostasis as a key factor in activity-dependent synaptic connectivity.

## Introduction

Understanding the rules governing synaptic wiring during brain development and their underlying molecular mechanisms ^1,2^, remains a major challenge for neuroscience. Neurons face unique challenges due to their complex shape, where local decisions on axonal connectivity are made distantly – both in time and space – from gene regulatory events in the cell body. While developmental transcriptional control of wiring molecules such as cell surface receptors is well accepted, it is not clear whether this also applies to fundamental cell biological processes that also act locally and are essential for circuit wiring, such as protein degradation or mitochondrial homeostasis. If this were the case, it would imply the existence of genetically encoded mechanisms of gene expression that regulate of local organelle biology during neuronal circuit development.

Several types of molecular mechanisms have been identified as necessary for the specificity of circuit wiring diagrams from the specification of neuronal identity driven by stereotyped expression of temporal transcription factors in neuronal nuclei ^3–5^, through neurite navigation via differential cell-surface proteins guidance ^6, 7^, developmental sorting and competition algorithms ^8, 9^ to partner selection and synaptogenesis ^10^, influenced by local cell biological processes such as autophagic degradation and control of filopodial dynamics ^11, 12^. Much remains to be learned about how these different mechanisms are integrated to produce highly stereotyped, yet individually variable, neuronal circuits ^13, 14^. Recent work in *Drosophila* and mouse have demonstrated that temporally expressed transcription factors play critical roles in axon targeting and circuit connectivity ^15–18^. Clearly, these temporally controlled regulatory events must regulate the expression of genes that then coordinate local events during the establishment of neuronal connectivity. Yet what these gene expression programs are and how they contribute to events such as axonal growth and the synapse formation remains to be elucidated.

Neuronal metabolism needs to be highly dynamic, requiring the development of appropriate maintenance mechanisms and leading to the constant rejuvenation of cellular organelles ^19^. Various local homeostatic processes were shown to be involved in neuronal circuit formation, including actin and microtubules turnover, autophagy, and mitostasis^19–21^. A crucial and highly dynamic organelle within the neuron are mitochondria, which are present both in the cell soma and distributed through the cellular periphery, including axons and dendrites. Mitochondrial function is tightly regulated by their structural dynamics, and they are capable to form distinct subpopulations within the neuronal compartments in response to specific cellular demands ^22–25^. In mature neurons, mitochondria are involved in synaptic transmission and plasticity regulation through local ATP supply and Ca^2+^ buffering ^24, 25^. During development, metabolic rewiring was shown to be a major player in quiescent neural stem cell activation and proliferation, and neurites outgrowth and branching are directly affected by the proper transport, dynamics, and anchoring of mitochondria ^26–28^.

Despite having their own genome, mitochondria depend crucially on genes encoded by the nuclear genome that are essential for mitochondrial biogenesis and function. In response to cellular energy and growth requirements mitochondrial mass and function are regulated by different sets of nuclear-encoded genes. Due to their importance in regulating processes such as energy production, calcium buffering, and cell survival, mitochondrial homeostasis is ensured by multiple systems of quality contro l^29^. Some of these systems ensure that damaged mitochondria are degraded by a process called mitophagy, resulting in continuous mitochondrial turnover and the renewal of the healthy mitochondrial pool. Proper maintenance of the axonal mitochondrial pool is now widely thought to be important for brain health and preventing diseases, such as neurodegeneration ^30^. Mitochondrial dynamics are important for axonal targeting, synaptogenesis, and circuit maintenance. Their metabolic activity sets the species-specific tempo of neuronal development, and is proposed to affect the dynamics of transcription during brain development ^31^.

The fly visual system, with its well-described developmental programs of cell patterning and circuit wiring, has been a powerful tool to dissect the molecular mechanisms of neuronal circuit formation and to extract the fundamental rules that these mechanisms subserve. Both retinal neurons and higher order visual system neurons, such as the Dorsal Cluster Neurons (DCNs/LC14s) have been used to identify genetic hierarchies that lead to the emergence of neuronal circuit wiring specificity ^32, 33^. Using the DCNs as a model system we report the discovery of a conserved temporally expressed transcription factor called *dTzap (drosophila TZAP)*, required for axonal growth and activity-dependent circuit connectivity. We show that dTzap is a transcriptional regulator of key mitochondrial homeostasis genes and loss of dTzap leads to structural and functional defects in axonal mitochondria. These defects can be rescued upon re-expression of its target genes, notably the Parkinson’s disease gene Pink1, demonstrating a causal link. Thus, developmental transcriptional regulation of mitochondrial homeostasis is required for neuronal circuit wiring.

## RESULTS

### DCNs transcriptome analysis

DCNs are a cluster of 22-68 cells per hemisphere marked by their dorsal location and expression of the transcription factor Atonal (Ato). They are divided into 2 subtypes based on their axonal target choice: a minority of DCNs innervate the contralateral medulla neuropile and are called M-DCNs (Figure 1A), while the majority innervate the lobula neuropile and are called L-DCNs ^9^ (Figure 1A’). As a rare and highly variable cell type, DCNs tend to not be highly represented in single-cell RNAseq data ^34^ and cannot be unambiguously distinguished from other Ato+ neurons, such as the Ventral Cluster Neurons (VCNs) ^35^. To specifically profile the DCN transcriptome, we used Laser Micro Dissection, followed by bulk next-generation RNA sequencing (RNA-SMARTseq2) ^35^ of isolated DCN clusters. We generated a range of microdissected DCNs samples and randomly cut central brain regions of the same animals as control samples (Figure 1B). After MDS clustering analysis of the top 1000 variable genes, we retained 6 bulk samples of DCNs and 2 control samples for further investigation (Supplementary File 1, tab1). The molecular identity of isolated DCN clusters was confirmed by strong expression of the marker gene *atonal* ^35^, which was missing in control samples. Housekeeping genes were expressed at similar levels (Figure 1C). We used a range of threshold parameters (logFC > 2, adjusted p-value ≤ 0,01, a mean base number ≥ 100) and manual analysis of available gene annotations to filter the list of candidate genes for further analysis (Figure 1D, Supplementary File 1, tabs1,2).

**Figure 1.**
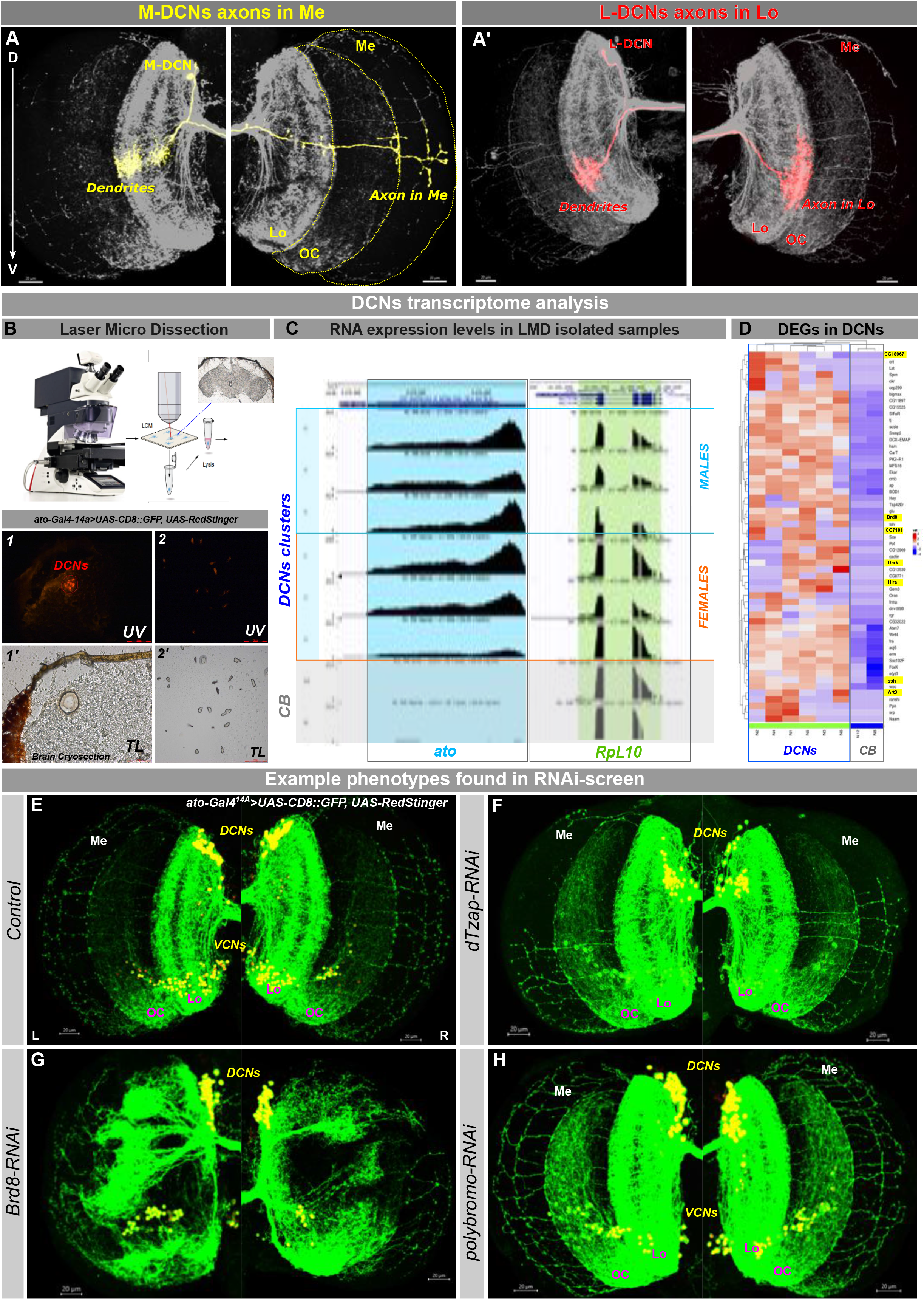
DCNs transcriptome analysis with subsequent RNAi screen of 74 high confidence genes. **A, A’** – Multi Color Flip-Out (MCFO) staining of single DCNs innervating either the medulla (M-DCN, **A**) or the lobula (L-DCN, **A’**). **B** – Laser Micro Dissection of DCN clusters from brain cryosections of flies *ato-Gal4^14A^; UAS-CD8::GFP, UAS-RedStinger*. 1,1’-RedStinger fluorescent DCNs before (1, UV light) and after (1’, transmitted light) dissection. 2,2’ - sample with dissected DCNs clusters in the UV (2) and transmitted (2’) light. Total RNA was extracted using Ambion RNAqueous Micro Kit and libraries were generated using SMART-seq2 Kit and Nextera Tagmentation Kit. NGS NextSeq 500 High Output Kit (400 million reads of approximately 25 million reads/sample coverage was used for sequencing). **C** - example of the marker gene *atonal* high-level expression in DCN samples compared to random central brain regions used as a control, with the equal expression of housekeeping genes. **D** - Heatmap of top-67 genes expressed in DCNs **E-H** – RNAi-screen of 74 DCN genes revealed 10 candidates causing significant phenotypic changes upon DCN-specific developmental downregulation. Lo – lobula, OC – optic chiasm, Me – medulla, DCNs – Dorsal cluster neurons, VCNs – ventral cluster neurons. Scale bar – 20µm.

We reasoned that some of the DCN-expressed genes might encode factors required for regulation of target genes that in turn control local axonal mechanisms during circuit assembly. If so, it should be possible to identify such factors through a genetic screen for DCN connectivity defects. To test this idea, we performed an RNAi-screen of 74 significantly enriched genes in DCNs and analyzed changes in DCN wiring pattern. As a first filter, we scored the number of DCN medulla axons and set a phenotypic threshold of more than 30% change in the mean number of M-DCN axons with a p-value lower than 0.01 compared to control flies (Supplementary Figure S1). This resulted in 10 candidate genes whose downregulation decreased, increased, or completely impaired medulla innervation, some in a sexually dimorphic manner (Figure 1E-H). Among those were transcription factors (*CG7101*), genes involved in chromatin remodeling (*Brd8, polybromo, Hira*), cytoskeleton organization (*Cnb*), and actin filament dynamics (*ssh*). Here we report the functional analysis of the previously uncharacterized role of the transcription factor encoded by *CG7101* in neural circuit development.

### CG7101/dTzap downregulation causes DCNs Medulla innervation loss

We identified *CG7101*, a gene encoding a zinc-finger transcription factor homologous to mammalian *TZAP/ZBTB48*, known for its role in telomere length control and mainly studied in the context of cancer biology ^36, 37^. We therefore rename *CG7101 as dTzap* (*Drosophila* TZAP). Using single-cell RNA sequencing data ^38^ we found that *dTzap* mRNA is moderately expressed in developing DCNs at 15 hours after puparium formation (P15) and is then progressively downregulated throughout pupal development, before being expressed at very low levels in adult flies (Supplementary Figure S2A). This is also true across neuronal cell types in the developing brain (Supplementary Figure S2B). Using an endogenously GFP-tagged version to examine the developmental expression profile of dTzap protein in DCNs, we found expression starting at P30, followed by downregulation at P50 and very low levels of expression in DCN cells in adults (Figure 2A, A’-E,E’). Next, we confirmed the loss of medulla axons phenotype observed in the initial screen using two independent RNAi lines *(#100127*, *#27849*) and CRISPR/Cas9 approach ^39^, by expressing a dTzap gRNA and Cas9 specifically in DCNs (Figure 2F-I). This phenotype was not degenerative, as we did not detect differences between M-DCN axons number in young and old flies in both control and dTzap-RNAi background (Supplementary Figure S2C). Thus, dTzap, encoded by *CG7101,* is a developmentally regulated transcription factor required for neuronal circuit development.

**Figure 2.**
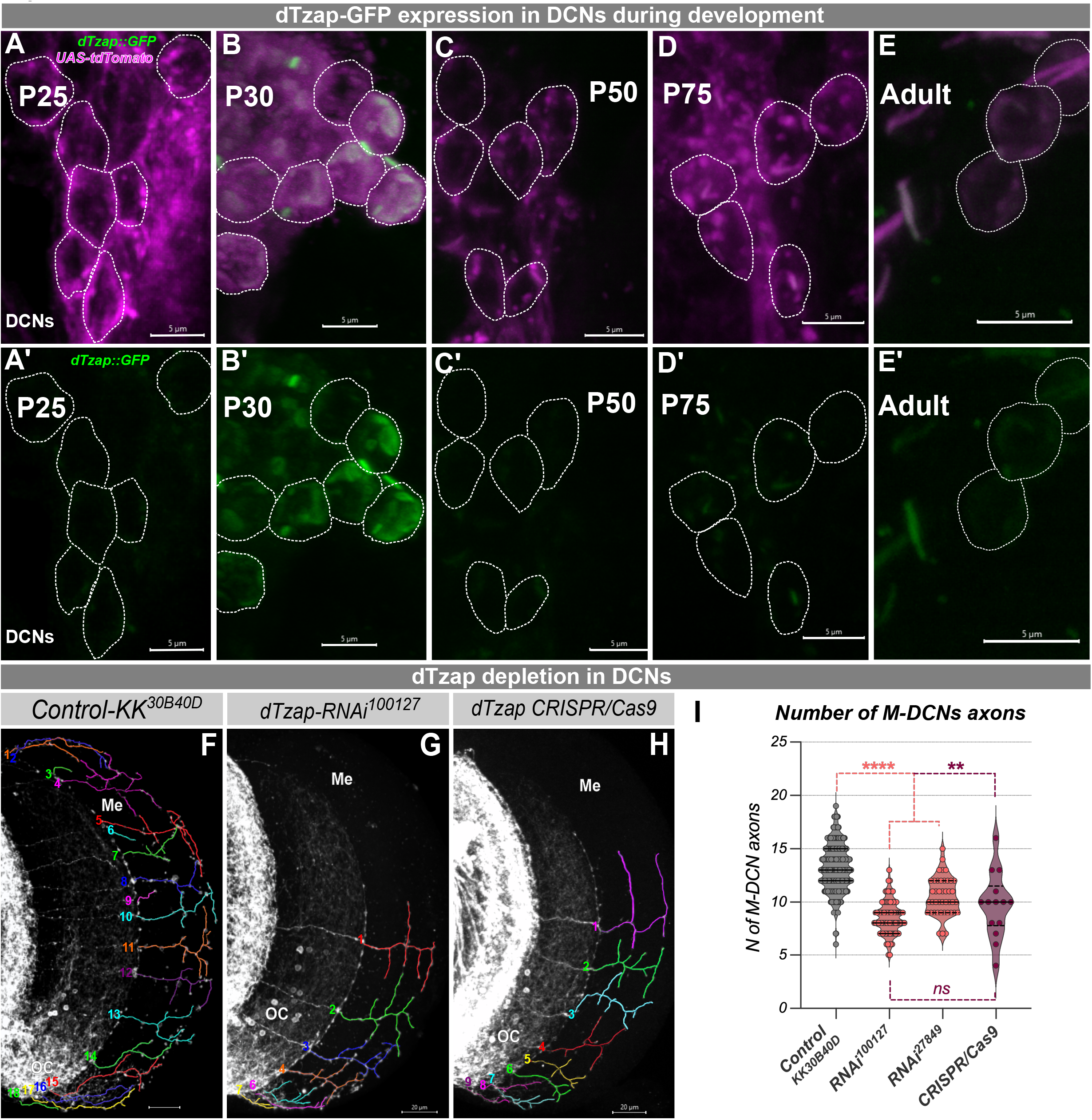
CG7101/dTzap is temporally regulated and its downregulation causes significant reduction of DCN axons in medulla. **A-E’** – temporal expression pattern of the tagged version of dTzap protein (*67655^BDSC^* combined with *ato-Gal4^14A^,UAS-CD4::tdTomato* transgenes) during development DCNs shows highest peak of dTzap protein expression at 30 hours after puparium formation (P30) with subsequent decrease, scale bar – 5µm. **F-I** – downregulation of *CG7101/dTzap* using independent RNAi lines (*100127^VDRC^* and *27849^VDRC^*) and CRISPR/Cas9 approach (*dTzap_sgRNA^341305^* and *UAS-U^M^-Cas9^34007^* from Heidelberg CFD CRISPR library) in DCNs (driving with *ato-Gal4^14A^*) leads to a significant decrease in the number of DCNs axons in medulla (I). OC – optic chiasm, Me – medulla, scale bar – 20µm. We examined 126 optic lobes of control group flies (males and females), 83 of *RNAi^100127^*, 42 of *RNAi^27849^* and 14 of CRISPR/Cas9. Statistical analysis was done using Kruskal-Wallis test with the Dunn corrections for multiple comparisons. *\* p<0.05, ** p*≤*0.01, *** p*≤*0.001, **** p*≤*0.0001, ns* – not significant.

### dTzap/TZAP binds to and regulates mitochondria homeostasis genes

Mammalian TZAP/ZBTB48 is a zinc finger transcriptional factor, which is composed of an N-terminal BTB/POZ domain and eleven adjacent C2H2-type zinc fingers (Znf1-11) at its C-terminus ^40^. As dTzap and human TZAP/ZBTB48 proteins show 33% similarity and 24% identity mostly in the zinc finger domains (Supplementary Figure 3A). We therefore wondered whether their function is conserved. As telomere length control is regulated differently in flies and humans, we reasoned that the role of dTzap in neuronal wiring is due to the regulation of other processes. It was reported that human TZAP activates the expression of the *Mitochondria Fission Process 1 gene (MTFP1)*, suggesting that human TZAP may be a regulator of mitochondrial homeostasis genes. To test whether fly dTzap and human TZAP might regulate other mitochondrial homeostasis genes, we analyzed available CG7101/dTzap ChIPseq data from the ModERN project ^41^ and TZAP ChIPseq data from human cell lines generated in previous studies ^42^. Our analyses reveal that, in addition to *MTFP1*, TZAP also shows binding peaks in genes encoding other mitochondrial regulatory proteins including *DNM1L, PINK1, ULK1*, *NRF1,* and *MFN2* (Figure 3A). Similarly, we found that dTzap has binding peaks to the promoter regions of the fly homologs of these human genes involved in mitochondria homeostasis, namely *CG7772/dMtfp1, Drp1*, *Pink1, Atg1, ewg, and Marf1* (Figure 3B). To test if TZAP/dTzap regulates these genes, we examined their relative mRNA levels upon TZAP downregulation in mammalian COS7 cells using si-RNA or post-mitotic pan-neuronal knock-out of *dTzap* in fly brains (*snyb-Gal4*>*dTzap_sgRNA,UAS-U^M^-Cas9^340007^*). This resulted in the loss of 80%-90% of TZAP/dTzap expression and significant downregulation of the 6 mitochondrial homeostasis genes tested in mammalian cells and in fly neurons (Figure 3C, D). Interestingly, overexpression of dTzap did not upregulate its target genes (Figure 3E), suggesting that dTzap is a developmentally regulated transcription factor required for the initial activation or upregulation of mitochondrial homeostasis genes in neurons.

**Figure 3.**
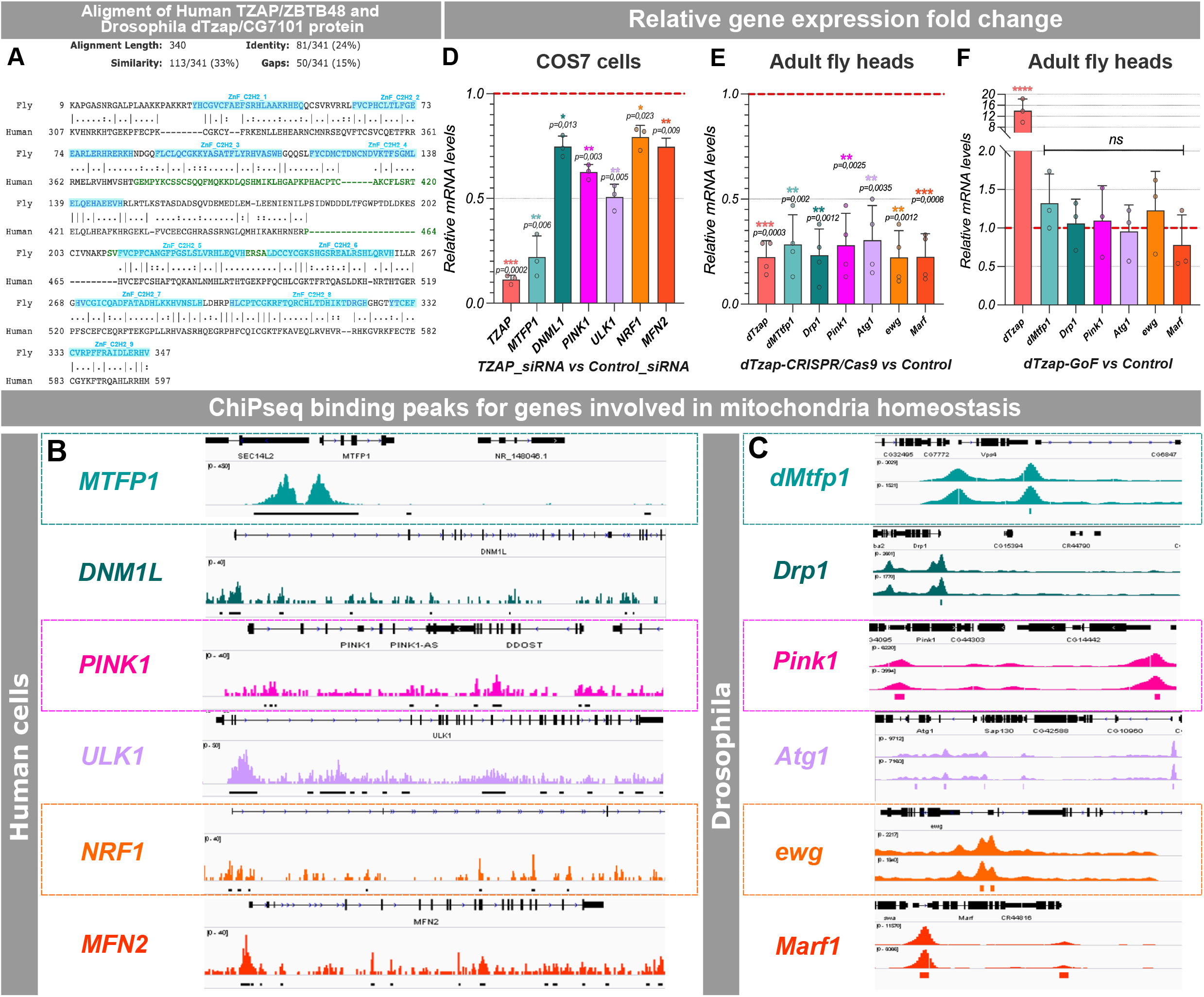
dTzap/TZAP is a conserved transcription factor that binds to and regulates nuclear genes involved in mitochondria homeostasis in flies and human cells. **A, B** – Analysis of available ChiPseq datasets from mammalian TZAP (A) and fly dTzap (B) in both cases showed peaks in a range of nuclear genes involved in mitochondria homeostasis at promoter regions. **C-E** – qRT-PCR measurement of relative mRNA levels of TZAP/dTzap target genes shows significant decrease in case of its downregulation with *Tzap_siRNA* in COS7 cells (C) and *nsyb-Gal4>dTzap_sgRNA;UAS-U^M^-Cas9^340007^* in flies (D). Overexpression of dTzap (*nsyb-Gal4>UAS-dTzap^F001205^)* had no effect on downstream genes expression (E). ΔCt values of the qRT-PCR experiment were compared using one-way ANOVA with Dunn correction for multiple comparisons. *\* p<0.05, ** p*≤*0.01, *** p*≤*0.001, **** p*≤*0.0001, ns* – not significant.

### dTzap acts as a conserved regulator of the mitochondria morphology

Mitochondria are crucial organelles for energy production, regulation of cell signaling, and amplification of apoptosis, and their function is tightly associated with their morphology ^43, 44^. We expressed GFP-tagged mitochondrial localization sequence (*UAS-mito-GFP*) in *Drosophila* DCNs and examined mitochondrial morphology in axonal terminals. Downregulation of *dTzap* in DCNs led to a significant increase in mitochondrial size in *dTzap-RNAi^1001^*^27^ and *dTzap_sgRNA, UAS-UM-Cas9* expressing flies (Figure 4A-F). More interestingly, we observed the same mitochondrial elongation in flies with developmental-specific *dTzap* downregulation, but not in flies with dTzap downregulation exclusively in adult flies (using *tub-Gal80^ts^* repressor, Figure 4B, B’’, D, D’, F). This observation confirms the requirement of dTzap during development for proper mitochondria morphology, which cannot be rescued by delayed expression in the adult.

**Figure 4.**
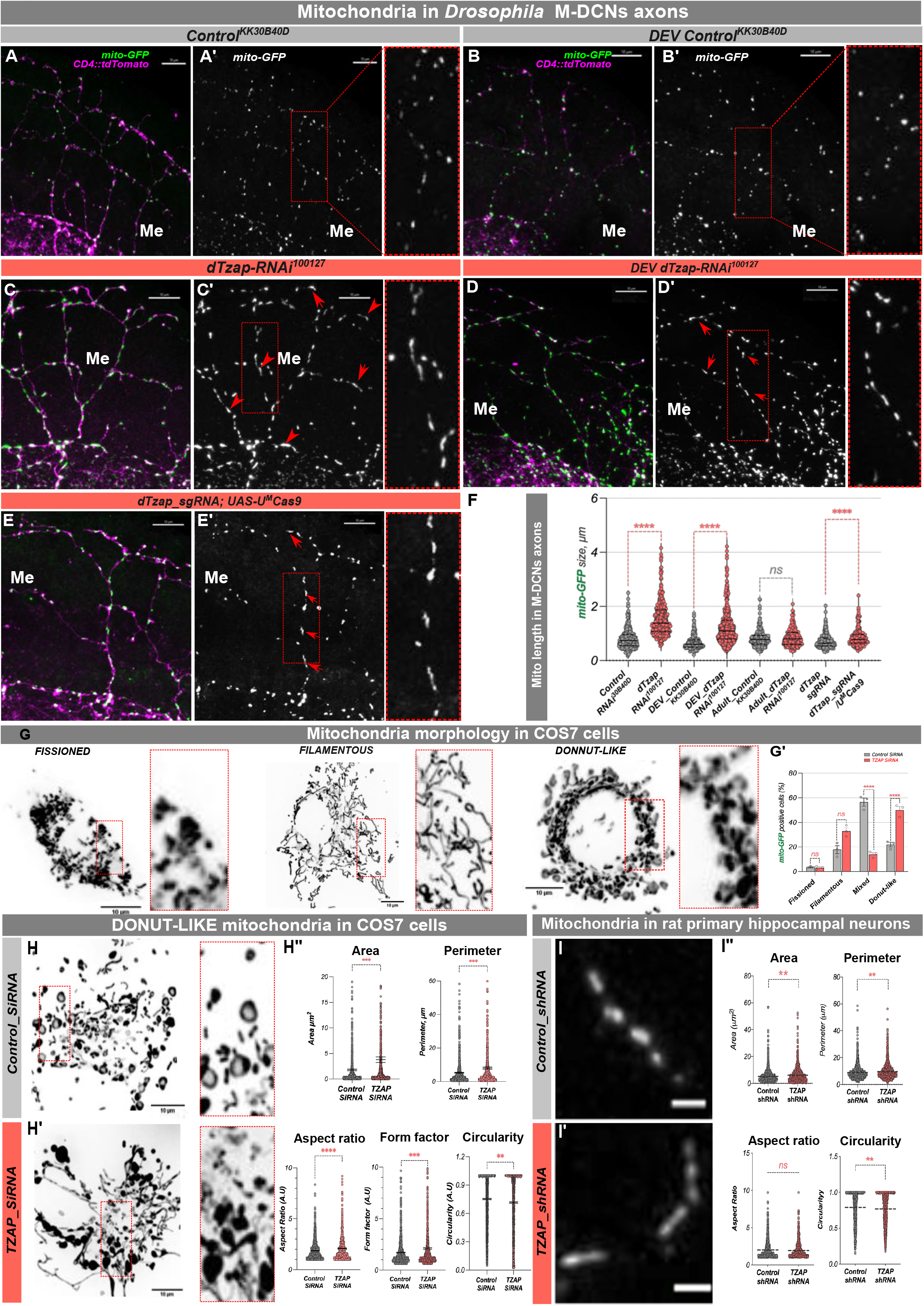
TZAP downregulation in Drosophila, rat hippocampal neurons and mammalian COS7 cells leads to defects in mitochondrial size and morphology. **A-E’** – Mitochondria in M-DCNs axonal terminals labeled with mito-GFP in Control condition (*w^1118^;UAS-mito::GFP/UAS-KK^30B40D^;ato-Gal4^14A^,UAS-CD4::tdTomato*) and dTzap downregulation using RNAi against *dTzap* (C-C’, *w^1118^;UAS-mito::GFP/UAS-dTzap^100127^;ato-Gal4^14A^,UAS-CD4::tdTomato*) and CRISPR/Cas9 system (E-E’, *w^1118^;UAS-mito::GFP/UAS-dTzap_gRNA;ato-Gal4^14A^,UAS-CD4::tdTomato/UAS-U^M^Cas9^340007^*). B, B’ – control for temporal dTzap downregulation during development (*UAS-KK^30B40D^/UAS-mito::GFP;ato-Gal4^14A^,UAS-CD4::tdTomato/tub-Gal80^ts^*). D, D’ – temporal dTzap downregulation during development (*dTzap-RNAi^100127^/UAS-mito::GFP;ato-Gal4^14A^,UAS-CD4::tdTomato/tub-Gal80^ts^)*. **F** – Mitochondria size was manually measured using slice mode in IMARIS through the Z-stack of a confocal image of one/two axons in the optic chiasm and medulla region in 4 different brains of each gender, n = ∼300 mitochondria per genotypes. Scale bar – 10µm. **G-G’** – Analysis of mitochondrial morphology of *TZAP* downregulation (*TZAP siRNA*) and control (*Control siRNA*) in mammalian COS7 cells. We were counting the number of cells that were assigned into four main classes of dominant mitochondria shapes: fissioned, filamentous, mixed, and donut-like. Downregulation of TZAP in mammalian COS7 cells leads to a shift in the proportion of mitochondrial morphology towards filamentous and donut-like shapes (G’). **H-H’** – Severe cases representation of cells with predominant donut-like mitochondria in Control (H) condition, and TZAP-RNAi (H’) with a lot of dense blobs structured mitochondria. Scale bar – 10µm. H’’ – measurement of mitochondria morphology parameters in *Control* and *TZAP-RNAi* conditions indicates an increase in area, perimeter, aspect ratio, and decrease in circularity and form factor in case of *TZAP* downregulation. Mitochondria in mutant cells tend to be bigger and more elongated, suggesting impairments in fusion-fission events balance. **I-I’’** – Measurement of mitochondria morphology parameters calculated in cultured rat primary hippocampal neurons in control and *TZAP-RNAi* conditions indicates an increase in area, and perimeter and a decrease in circularity in case of *TZAP* downregulation, Scale bar 5µm, n = 1397 mitochondria analyzed for *Control_siRNA*, and 1218 – for *TZAP_siRNA* condition. Statistical analyses were performed using GraphPad Prism 8 software. Normal distribution was checked using the D’Agostino Pearson normality test before statistical comparisons. A nonparametric Mann-Whitney U test was used for mitochondria measurements in fly DCNs and rat cortical neurons. For COS7 cells data were analyzed by two-way ANOVA, with Tukey post hoc test for multiple comparisons. The sample size was standard and determined based on previous experience. *\* p<0.05, ** p*≤*0.01, *** p*≤*0.001, **** p*≤*0.0001, ns* – not significant.

Mitochondria are dynamic organelles, and their shape and size are precisely controlled by fusion and fission events, which should be balanced for maintaining overall mitochondrial morphology ^45, 46^. To investigate mitochondrial morphology at higher resolution we examined the effects of TZAP silencing in the COS7 cells expressing mito::GFP, which are well-suited for due to their large and flat cytoplasm and well-developed mitochondrial network. Under control conditions, most cells (60%) contain a mix of mitochondrial shapes, while 20% of the cells show predominantly fused filamentous mitochondria, and another 20% show “donut-like” mitochondria, indicative of cellular stress (Figure 4G, G’). Following transfection with TZAP siRNA there was a significant increase in the proportion of cells with donut-like mitochondria (from 22% to 50%), a tendency towards more cells with fused filamentous mitochondria (from 17% to 33%), and a concomitant decrease in the proportion of cells with mixed mitochondrial morphologies, compared to the control siRNA condition (Figure 4G, G’). In addition, in some severe cases within siRNA-TZAP transfected cells, we observed the formation of blob shape mitochondria (Figure 4H-H’’) indicating irreversible toxicity ^44^. Quantitative image analysis of mitochondrial morphology in individual cells with a predominantly donut-like mitochondrial network revealed an increase in the mean values of aspect ratio and form factor following *TZAP* silencing compared to control cells, two parameters reflecting the length and the degree of branching of the mitochondrial objects. In addition, the mean values of the perimeter and area of mitochondrial objects were increased, and circularity was decreased in cells exposed to TZAP siRNA compared to cells treated with Control siRNA (Figure 4H-H’’).

Next, we asked what the impact of TZAP loss on mammalian neuronal mitochondrial morphology might be. We expressed a mitochondria-localized Ca^2+^ sensor (mito^4x^-GCaMP6f)^47^ in sparsely transfected rat primary hippocampal neurons, which allowed us to identify single mitochondria in axons and evaluate their morphology using the baseline mito^4x^-GCaMP6f (Figure 4I-I’’). We thus evaluated the area, perimeter, circularity, and aspect ratio and found that *TZAP_shRNA* neurons presented larger mitochondria with a more elongated shape, confirming our observations in mammalian COS7 cells and *Drosophila* neurons.

Altogether, our results so far show that *dTzap/TZAP* is developmentally regulated transcription factor that is required to maintain normal mitochondrial size likely through the regulation of expression of several mitochondrial homeostasis genes.

### TZAP downregulation impairs mitochondrial Ca^2+^ uptake and presynaptic glutamate release

To evaluate whether reduced expression of mammalian *TZAP* may alter the ability of mitochondria to uptake Ca^2+^ during neuronal activity, we co-expressed mito^4x^-GCaMP6f with *TZAP shRNA* and quantified axonal mitochondrial Ca^2+^ uptake during neuronal activity induced by field stimulation of a train of 20 action potentials at 20Hz (Figure 5A). We observed that upon *TZAP* knock-down, a higher percentage of neurons failed to show any quantifiable response in comparison to controls (Figure 5B). Among those neurons responding, we observed significantly reduced peak mitochondrial Ca^2+^ response, yet mitochondrial Ca^2+^ extrusion remained unaltered (Figure 5C). To test whether TZAP expression can control presynaptic function in mammalian neurons, we expressed a surface-localized glutamate sensor, iGluSnFR3 in primary hippocampal neurons, at the presence or absence of an shRNA against *TZAP*. iGluSnFR3 exhibits fast non-saturating activation dynamics and reports presynaptic glutamate release with specificity ^50^. Single action potential firing generated robust presynaptic iGluSnFR3 signals (Figure 5D; black trace, Figure 5E) as previously reported ^48^. However, we observed that glutamate release was reduced by ∼50% in TZAPshRNA-expressing neurons, indicating that in absence of TZAP presynaptic neurotransmitter release is impaired (Figure 5E’). Thus, the absence of TZAP in hippocampal neurons results in aberrant mitochondrial calcium dynamics and reduced neurotransmitter release.

**Figure 5.**
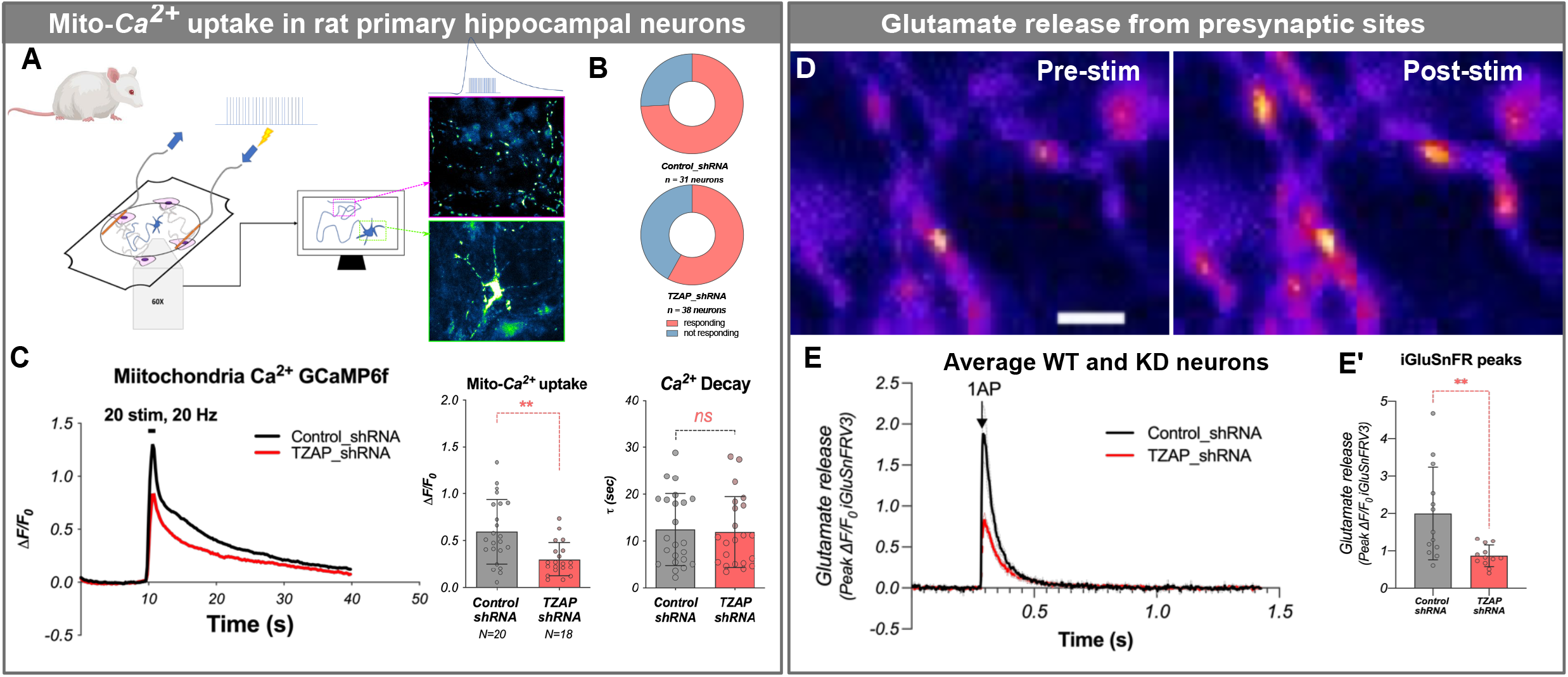
TZAP downregulation decreases mitochondrial Ca2+ uptake and presynaptic neurotransmitter release in primary hippocampal neurons. **A**-General scheme of the experiment. Primary neurons derived from rat hippocampus were cultured and sparsely co-transfected with vectors, carrying GCaMP6f and *Control_shRNA* or *TZAP_shRNA* transgenes. **B** - Percentage of responding and nonresponding cells in *Control_shRNA* and *TZAP_shRNA* conditions. **C** – Representation of axonal mito-Ca^2+^ responses to 20AP (20Hz, 1ms) stimulation, calculation of mito-Ca^2+^ uptake and decay in mature neurons (DIV 14-21), *n* = 20 for *Control_shRNA* neurons, and 18 for *TZAP_shRNA*. **D** - Representative pseudo color images of glutamate release from presynaptic sites using iGluSnFR sensor. **E** - Representative average presynaptic responses in Control and *TZAP-RNAi* conditions. **E’** - Calculations of iGluSnFR fluorescence peaks (Δ*F/F_0_*) in both conditions, *n* = 13 for *Control_shRNA* neurons, and 12 for *TZAP_shRNA*. Scale bar – 5µm. Statistical analysis was performed with GraphPad Prism 8 using the nonparametric Mann-Whitney U test. *\* p<0.05, ** p*≤*0.01, *** p*≤*0.001, **** p*≤*0.0001, ns* – not significant.

### Loss of dTzap leads to loss of activity-dependent synaptic connectivity and corresponding behavioral defects

To examine the role of TZAP/dTzap in synaptic connectivity and neuronal circuit function *in vivo* we investigated synapse formation in *Drosophila* DCN axons. Although loss of dTzap reduces the number of DCN medulla targeting axons, 40%-50% of the expected DCN axons still target the medulla and appear to have normal terminal morphology. Furthermore, *in vivo* pyruvate measurement ^48^ revealed that knock-down of dTzap did not lead to defects in pyruvate accumulation upon blockade of mitochondrial respiratory chain, indicating no major defects in mitochondrial metabolism (Supplementary Figure S4). Next, we examined the impact of dTzap loss on DCN circuit connectivity and function. We began by examining the presynaptic markers Brp, Syd1, and Nrx1, by expression of GFP-tagged versions and found no qualitative or quantitative differences between *dTzap-RNAi* and control DCNs (Supplementary Figure S5A-C”). Thus, loss of dTzap does not appear to impair the formation of presynaptic sites. As we observed reduced neurotransmitter release from presynaptic sites in hippocampal neurons upon Tzap depletion, we examined synaptic vesicles release in M-DCN axonal terminals by performing live imaging of a dual-tagged pH sensitive sensor pHluorin, fused to stable mCherry fluorophore and Synaptotagmin (Syt1-mCherry::pHluorin) ^49^. This sensor allows the localization of presynaptic release sites with the pH resistant mCherry, while simultaneously permitting quantification of synaptic vesicle exocytosis with the pH-sensitive GFP moiety, which will fluoresce upon vesicle release exposing GFP to the neutral pH of the extracellular environment. We observed a dramatic decrease in GFP pHluorin signal normalized to the mCherry signal in case of dTzap downregulation suggesting significant reduction in synaptic vesicle release in DCN axons after dTzap depletion (Figure 6A-A’’, B-B’’, C). To test whether this loss of activity impaired neuronal circuit connectivity, we used the anterograde transsynaptic tracing approach TransTango ^50^ and quantified the number of positively labeled postsynaptic cells in the medulla (Figure 6D, E,F,F’’, magenta). This revealed a ∼10-fold loss of postsynaptic connectivity with target neurons upon *dTzap* downregulation with both RNAi and CRISPR/Cas9 approaches (Figure 6D-E’’, G, Supplementary Figure S5D-D’, G, white arrowheads). This reduction did not lead to a change in targeting specificity, because the few cells that did label in dTzap loss of function conditions corresponded to the previously identified postsynaptic targets of DCNs, like Lawf1/2, Tm2/21, DCN, and Dm3/6 neurons ^51^. Thus, dTzap is required for presynaptic vesicle release and neuronal circuit connectivity.

**Figure 6.**
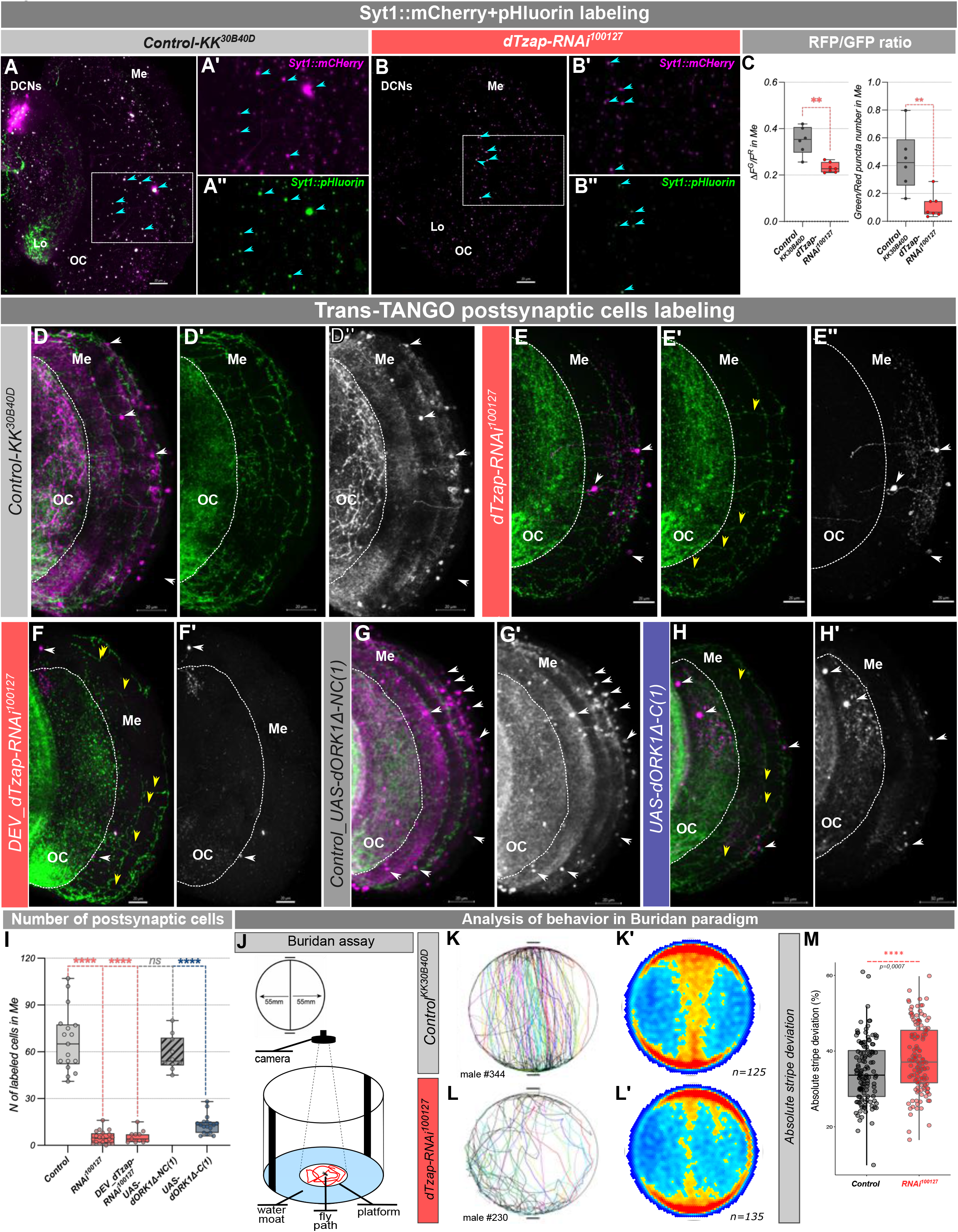
dTzap downregulation impairs synaptic vesicle release and circuit connectivity. **A-B”** – Live imaging of neurotransmitter release using UAS-Syt1-mCherry::pHluorin pH sensitive fluorescence probe in control (A,A’) and *dTzap-RNAi^100127^* (B,B’) condition. **C** – Ratio of green (pHluorin, cyan arrowheads) to red (mCherry) fluorescence and number of puncta in control (n=6 flies) and dTzap downregulation condition (n=7 flies). **D-E”, G** – TransTango anterograde postsynaptic labeling revealed that *dTzap* downregulation, with an expression of *UAS-dTzap-RNAi^100127^* (n=20 optic lobes) *and dTzap_gRNA; UAS-U^M^-Cas9* (Supplemental figure S5, n=25 optic lobes) causes a dramatic loss of synaptic connectivity of DCNs with their postsynaptic partners compared to control (n=17 optic lobes), despite the visible M-DCN axons (yellow arrowheads). **F’’, F’’’** – Silencing of DCNs by overexpression of conductive form of potassium channel dORK (*UAS-dORK1*Δ*-C(1),* n=18 optic lobes) showed a comparable reduction of postsynaptic cell labeling. F, F’ – overexpression of the nonconductive form of dORK (*UAS-dORK1*Δ*-NC(1)*, n=10 optic lobes) used as an additional control, showed no effect on TransTango labeling. DCNs expressing myrGFP on the cellular membrane are shown in green (D, D’, E, E’, F, F’’) and postsynaptic cells in magenta (merged picture) and grayscale (D, D’’, E, E’’, F-F’’’). Postsynaptic connectivity was analyzed by manually counting cell bodies in medulla (white arrowheads, G) labeled with dsRed fluorescence (magenta, merged picture – D, E, F, and grayscale – D”, E”, F”), from their cell bodies, including all cells with weak or strong labeling to reveal all potential connections. Scale bar 20µm. We examined the object orientation behavior of flies with dTzap downregulation in DCNs in Buridan assay (H) and found that it was altered compared to the control. I – **J** – Representation of individual fly trajectory during recording. I’, J’ – heat map of arena occupancy at the population level. We examined individual behavior of 125 control flies and 135 *RNAi^100127^* flies (together males and females in equal proportion). **K** – Absolute stripe deviation measurement, showing that Control flies (*W+; UAS-KK^40D30B^;ato-Gal4^14A^,UAS-CD4::GFP*) normally walk more straight (represented in lover ASD value) than flies with dTzap downregulation (*W+;UAS-dTzap-RNAi;ato-Gal4^14A^,UAS-CD4::GFP)*. Me – medulla, Lo – lobula, OC – optic chiasm. Statistical analysis was done using Ordinary one-way ANOVA with the Kruskal-Wallis test for the behavioral experiment an unpaired Student’s t-test with Welch’s correction for synaptic marker counting, and one-way ANOVA with Bonferroni corrections for multiple comparisons in trans-TANGO labeling analysis. *\* p<0.05, ** p*≤*0.01, *** p*≤*0.001, **** p*≤*0.0001, ns* – not significant.

The loss of TransTango signal was somewhat surprising because TransTango is not thought to be an activity-dependent method, although this has never been specifically tested to our knowledge. To test whether this is the case, we blocked DCN activity using three different silencers: inhibiting synaptic vesicle exocytosis via Tetanus toxin (*UAS-TNT*) ^52^, or suppressing electrical activity by expression of potassium channels (Kir2.1 or dORK) that are open at resting membrane potential which results in increased potassium efflux and membrane hyperpolarization setting resting membrane potential below the threshold required to fire action potentials ^53^. In all cases, we saw the same drastic decrease of postsynaptic labeling observed in the *dTzap* downregulation condition (Figure 6F-G, Supplementary Figure S5E-G’). Importantly, expression of the nonconductive form of dORK as a control ^54^ showed postsynaptic labeling comparable to control flies (Figure 6F-F’, G).

To ascertain that the loss of TransTango postsynaptic labeling reflects loss of functional connectivity, we tested DCN behavioral function in dTzap loss of function flies. It was previously shown that DCN medulla targeting is required for object orientation behavior of freely walking flies, measured by absolute stripe deviation in the Buridan paradigm assay ^55^. Briefly, flies walk freely between two opposite and identical high-contrast stripes which they are unable to reach. The width of the path they trace during these back-and-forth walks is known as absolute stripe deviation (Figure 6H). We have previously shown that this parameter depends on DCN function and correlates with the degree of left-right asymmetry in medulla DCNs wiring ^14^. Silencing of DCN activity results in a significant increase of absolute stripe deviation at the population level because flies tend to walk across the entire arena and along the edges (Figure 6I,I’-J,J’). It also results in loss of the correlation between DCN connectivity and behavior in individual flies. Thus, absolute stripe deviation provides a quantitative parameter to measure DCN circuit function *in vivo*. We measured absolute stripe deviation in control and DCN-specific dTzap loss function conditions and observed a loss of the correlation between wiring and behavior in individuals (Supplementary Figure S6A-E) and an increase in absolute stripe deviation (Figure 6I-K) quantitatively similar to that observed upon UAS-TNT expression in DCNs ^14^.

Taken together, our data show that dTzap is a developmental transcription factor required for the establishment of activity-dependent synaptic connectivity and circuit function.

### dTzap regulates mitochondrial morphology and circuit connectivity via Pink1

dTzap regulates several genes involved in mitochondrial dynamics and homeostasis. To test which target genes are involved in what aspects of the *dTzap* phenotype, we downregulated five of them independently (*dMtfp1/CG7772, Drp1, ewg, Atg1, and Pink1*) using DCN-specific RNAi. Knockdown of *Pink1* and *Atg1*, but not other target genes, resulted in elongated axonal mitochondria (Figure 7A-D’, H), mimicking dTzap loss of function phenotype. Moreover, downregulating *Pink1*, but not *Atg1 or Drp1*, resulted in loss of TransTango labeling, reflecting compromised connectivity with postsynaptic targets (Figure 7E-G’, I). Thus, downregulation of *Pink1* phenocopied both mitochondrial morphology and postsynaptic labeling defects. We therefore asked if restoration of *Pink1* expression would at least partially rescue the phenotypes of dTzap loss of function. To this end, we combined *dTzap-RNAi* with *Pink1* overexpression in DCNs. This resulted in restoration of both mitochondrial size (Figure 8A-C”) and connectivity to postsynaptic cells back to control levels (Figure 8D-F”). The initial phenotype we observed upon *dTzap* RNAi was a DCN medulla axon targeting reduction. We did not observe a reduction in the number of medulla axons upon *Pink1* downregulation alone, suggesting that this phenotype maybe a compound phenotype of multiple target genes, However, to our surprise *Pink1* overexpression restored the number of DCN medulla axons to that observed in controls (Figure 8G-I). Together, these data suggest that *Pink1* is a key target gene of dTzap mediating its role in regulating neuronal circuit development via mitochondria homeostasis.

**Figure 7.**
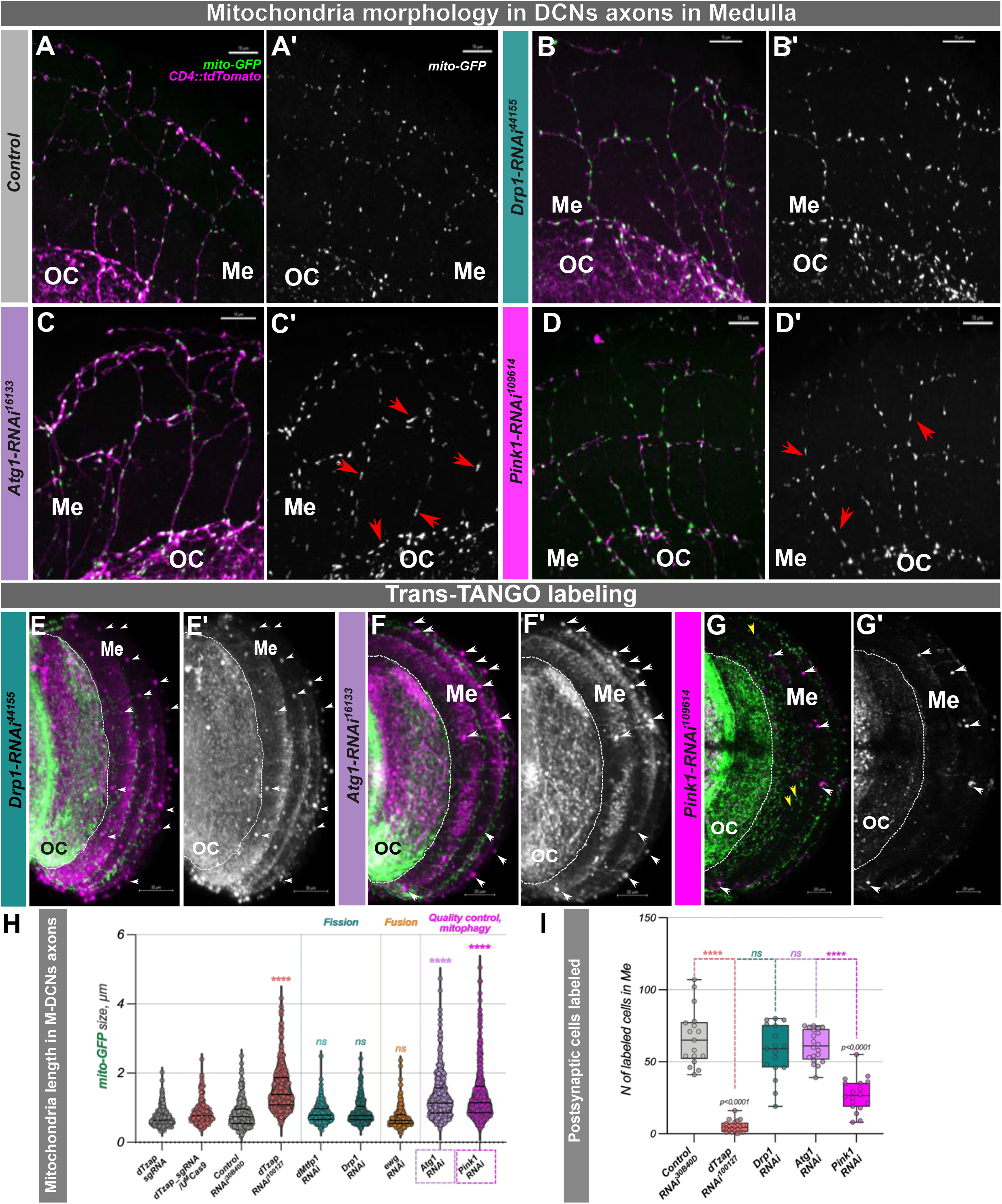
downregulation *of* dTzap target genes phenocopies dTzap loss of function. **A-H** – Mitochondria (labeled with mito-GFP) in DCNs axons in *dTzap* target genes downregulation flies compared to control (A). Shown are RNAi against *Drp1* (B, B’), *Atg1* (C, C’) and *Pink1* (D, D’). **H** – Quantification of mitochondria size in different genotypes measured through the Z-stack of a confocal image of 1-2 axons in the Optic chiasm and medulla region in at least 3-4 different brains of each gender. Scale bar – 10µm. **E-G’** – TransTango labeling of postsynaptic connectivity in *Drp1* (*Drp1-RNAi^44155^*, n=17 optic lobes), Atg*1* (*Atg1-RNAi^16133^*, n=20 optic lobes), and *Pink1* (*Pink1-RNAi^109614^,* n=18 optic lobes) downregulation in DCNs. **I** – The number of postsynaptic cells was calculated by manually counting cell bodies in medulla (white arrowheads) labeled with dsRed fluorescence (magenta and grayscale), from their cell bodies, including all cells with weak or strong labeling to reveal all potential connections. *Pink1* downregulation strongly reduced TransTango labelling around the M-DCN axons that are present (yellow arrowheads in G, G’). Scale bar – 20µm. Statistical analysis was done using the nonparametric Kruskal-Wallis test with Dunn correction for multiple comparisons and one-way ANOVA with Bonferroni correction for multiple comparisons respectively. *\* p<0.05, ** p*≤*0.01, *** p*≤*0.001, **** p*≤*0.0001, ns* – not significant.

**Figure 8.**
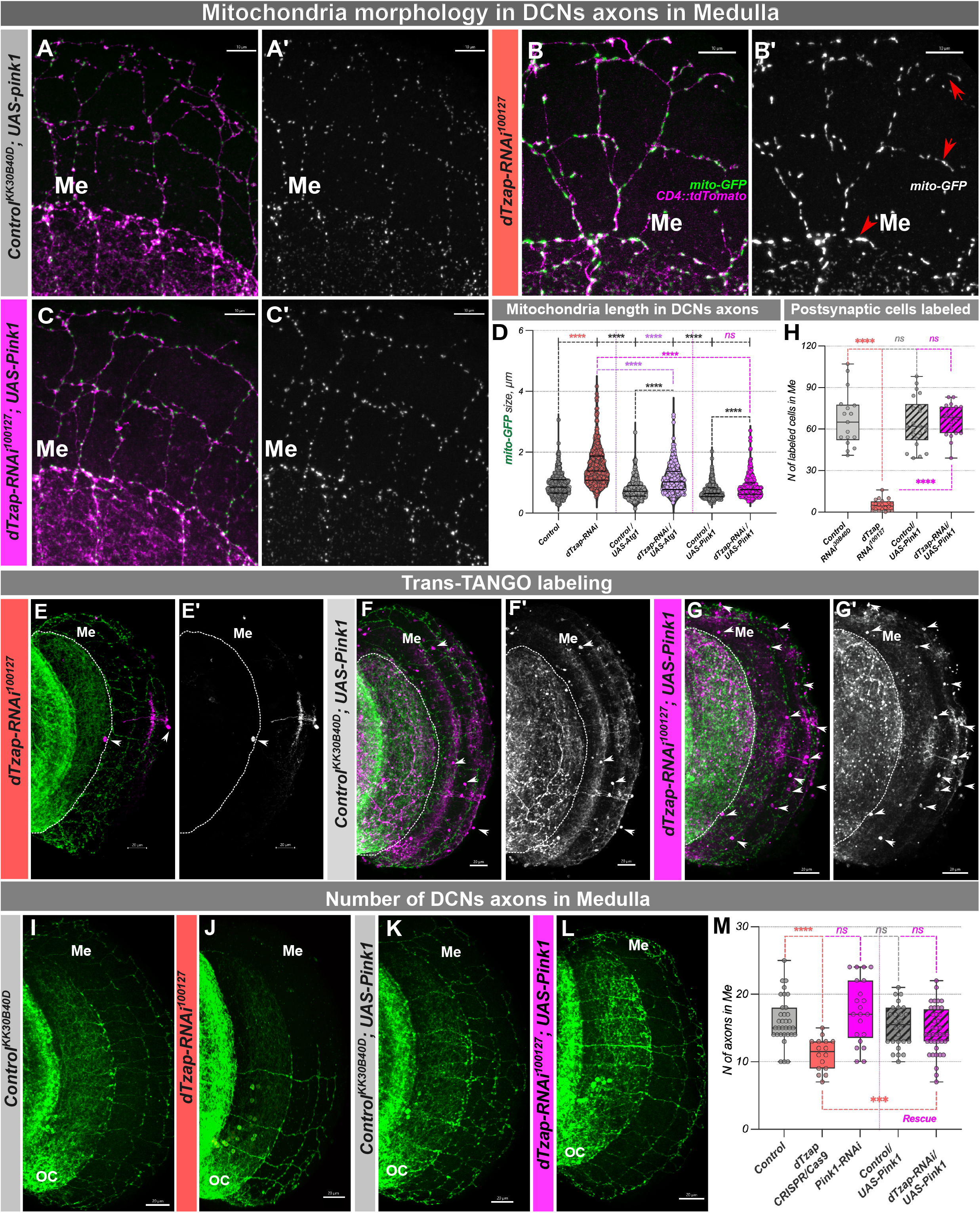
Pink1 expression rescues mitochondria size, postsynaptic labeling, and the number of DCN axons in the medulla caused by dTzap downregulation. **A-C’’**-Mitochondria size (labeled with mito-GFP) in DCNs axons was decreased to that observed in genetic controls (w^1118^;*UAS-mito::GFP/UAS-KK^40D30B^,ato-Gal4^14A^,UAS-CD4::tdTomato/UAS-Pink1*, A-A’) with *Pink1* overexpression (w^1118^;*UAS-mito::GFP/ UAS-dTzap-RNAi^100127^,ato-Gal4^14A^,UAS-CD4::tdTomato/UAS-Pink1*,C-C’). Mitochondria size was manually measured through the Z-stack of a confocal image of 1-2 axons in the Optic chiasm and medulla region in at least 3-4 different brains of each gender (C’’). Scale bar – 10µm. **D-F’** - anterograde transsynaptic labeling of postsynaptic partner cells using the TransTango approach showed complete rescue of connectivity in the case of *UAS-Pink1* expression in *dTzap-RNAi*^100127^ background (E’, F’, n=22 optic lobes for w^1118^;*UAS-mito::GFP/UAS-KK^40D30B^,ato-Gal4^14A^,UAS-CD4::tdTomato/UAS-Pink1* and 20 optic lobes for w^1118^;*UAS-mito::GFP/ UAS-dTzap-RNAi^100127^,ato-Gal4^14A^,UAS-CD4::tdTomato/UAS-Pink1* genotypes). The postsynaptic connectivity was calculated by manually counting cell bodies in medulla (white arrowheads) labeled with dsRed fluorescence (magenta and grayscale) (I), from their cell bodies, including all cells with weak or strong labeling to reveal all potential connections. Scale bar – 20µm. **G-H’** – Number of DCN axons reaching the medulla in adult flies. *Pink1* overexpression (*ato-Gal4^14A^>UAS-dTzap-RNAi^100127^,UAS-Pink1*) could rescue the number of M-DCNs axons decrease, caused by *dTzap* downregulation (I). Measurements were done equally in flies of both genders. Me - medulla. Scale bar - 20μm. Statistical analysis was done using one-way ANOVA with Dunnett, and Bonferroni corrections for multiple comparisons in mitochondria length comparison. *\* p<0.05, ** p*≤*0.01, *** p*≤*0.001, **** p*≤*0.0001, ns* – not significant.

In summary, dTzap is a conserved regulator of neuronal mitochondrial homeostasis genes required for the development of functional neuronal circuits.

## DISCUSSION

Deciphering the emergence of synaptic connectivity patterns requires understanding the mechanistic relationship between the regulation of gene expression in the neuronal nucleus, and local cell biological processes in distant neurites. To tackle this question, we carried out a cell-type specific transcriptomic and genetic screen in higher-order neurons of the *Drosophila* visual system called DCNs. Because the DCNs are an example of a rare cell type in the fly brain, it was challenging to ascertain their full and specific transcriptome from single-cell RNAseq datasets where typically a minority of DCN cells are represented. Similarly, other techniques, like TRAP (Translating Ribosome Affinity Purification) or TaDa (Targeted expression of the DamID) rely on using a cell-type specific Gal4 driver, not expressed anywhere else. To circumvent these limitations, we used Laser Micro Dissection followed by subsequent bulk next-generation RNA sequencing of isolated DCN clusters. With this approach, we were able to recover the expression of 8726 genes and reveal genes with low to medium expression levels, which could otherwise be easily missed. Although we determined the adult DCN transcriptome, an RNAi screen of 74 genes in postmitotic DCNs showed that several encoded nuclear factors that caused developmental connectivity phenotypes, including transcription factors like *Sox102F* and genes such as *Hira* ^56–58^ and *Brd8* ^59^, which encode a histone chaperone and a member of the Tip60 chromatin remodeling complex, respectively. This suggests that several levels of regulatory control are involved in circuit wiring in postmitotic neurons. Here we focused on the functional analysis of one of the genes identified in our screen, the transcription factor dTzap, the fly homolog of the human protein TZAP/ZBTB48, a BTB domain and Zinc Finger-containing (ZBTB) transcription factor.

TZAP, known for its role in negative regulation of telomere length and mainly studied in the context of cancer biology ^36, 37^, had not been studied in the nervous system. Since the mechanism of telomere maintenance in *Drosophila* and mammals are fundamentally different ^60, 61^, we reasoned that it was highly likely that dTzap’s potential role in circuit development occurred via the regulation of a different cellular process. Human TZAP/ZBTB48 had been proposed to be a transcriptional activator of a small set of target genes, including *Mitochondrial Fission Process 1* (*MTFP1*) ^42^. Mutation of the *Zbtb48* gene in mice caused a series of phenotypical manifestations, including abnormal bone structure and eye morphology defects in young adults, and impaired glucose tolerance and behavioral deficits at late adult stages ^62^, but the underlying mechanisms have not been studied. Thus, very little was known about the function of this highly conserved transcription factor in the brain.

We took advantage of available ChIP-seq data in human cells and in flies to ask what type of target genes TZAP/dTzap might regulate. This revealed a significant number of binds to genes well known for their requirement in mitochondrial homeostasis including the key mitochondrial quality control and Parkinson’s disease gene *PINK1/Pink1*, MTFP1/*dMtfp1* and the mitophagy and mitochondrial fission gene *DNM1L/Drp1* associated with developmental delay, ataxia and peripheral neuropathy ^63^. Mitochondria play a variety of vital roles. In addition to providing cellular energy, they also contribute to cellular calcium management providing rapid, post-stimulatory Ca^2+^ recovery by taking up massive amounts of calcium with subsequent gradual release into the cytosol ^64, 65^. Using a combination of *Drosophila*, COS7 cells and primary rat hippocampal neurons, we found that TZAP/dTzap plays a conserved role in regulating these mitochondrial homeostasis genes and that its loss leads to significant structural defects and reduced calcium import into mitochondria. Fluctuations in Ca^2+^ levels within mitochondria are known to impact their morphology and function with reduced calcium influx resulting in mitochondrial hyperfusion ^66^, consistent with our observations. We also observed strongly reduced glutamate release from rat hippocampal neurons. *In vivo*, loss of dTzap function led to drastic reduction in activity-dependent DCN postsynaptic connectivity and loss of DCN circuit function. Finally, on a technical note, our data also revealed that one of the most commonly used approaches for anterograde trans-synaptic tracing in *Drosophila*, TransTango, is in fact activity-dependent and this should be taken into account in interpreting TransTango data.

Knock-down of dTzap mitochondrial target genes, especially *Pink1*, phenocopied the *dTzap* loss of function phenotypes, supporting the causal role of their downregulation. Interestingly, and to some extent surprisingly, the restoration of Pink1 expression alone was able to rescue all the phenotypes associated with *dTzap* loss. This demonstrates that all the connectivity phenotypes caused by *dTzap* loss of function are likely due to mitochondrial defects and suggests that even when the level of expression of other genes is reduced, presence of sufficient amounts of *Pink1* is enough to ensure a level of mitochondrial function compatible with normal circuit development. Consistent with this, *Pink1* was shown to be a key component of the mitochondria quality control system initiating clearance of depolarized mitochondria by mitophagy ^67^. It is also shown to be an important regulator of cellular calcium turnover. Work using mice fibroblast cells has demonstrated that depletion of PINK1 leads to impairment in mitochondrial Ca^2+^ uptake after stimulation ^68^.

Although TZAP’s main function in the context of human cancer is thought to be in telomere length regulation, it has been observed that *TZAP* knockout in human cell lines led to abnormally clustered mitochondria ^42^. Interestingly, increasing evidence points to mutual crosstalk whereby telomere impairments lead to mitochondrial dysfunction and vice versa ^69^. It was recently shown that dysfunctional telomeres activate immune responses through mitochondrial TERRA-ZBP1 interaction, and act as a tumor-suppressor mechanism in human tissues ^70^. It would therefore be interesting to query whether telomere length regulation by mammalian TZAP and its transcription regulation of mitochondrial genes are coupled.

The main goal of our study was to identify developmental regulatory factors that couple gene regulation to local synaptogenic events during neuronal circuit development. Excitingly, this led to the discovery that a transcription factor that regulates mitochondrial homeostasis genes to facilitate activity-dependent connectivity is itself developmentally regulated. *dTzap* mRNA in DCNs peaks early during pupal development before being strongly downregulated until the adult stage where it is expressed at low levels. Importantly, the dTzap protein shows the same dynamics with an expected delay. This identifies dTzap as a novel postmitotic neuronal temporal transcription factor that couples developmental programs of gene expression to fundamental neuronal cell biology processes required for neuronal circuit formation.

## Acknowledgements

This work was supported by the Investissements d’Avenir program (ANR-10-IAIHU-06), Paris Brain Institute-ICM core funding, The Allen Distinguished Investigator Award (ADI12202), Roger De Spoelberch Prize, an NIH Brain Initiative RO1 grant (1R01NS121874-01), and ANR grant QuantSocInd (ANR-19-CE16-000) (to B.A.H.). We are grateful for use of and support from the following Paris Brain Institute’s core facilities and their staff: iGenseq for RNAseq, Histomics for histology and ICM.Quant for imaging. We thank Dr. Christian Lobsiger for help with Laser Microdissection, all members of the Hassan lab for helpful discussions and Dr. Nicolas Renier for comments on the manuscript.

## Author contributions

I.M. and B.A.H. conceived the study, designed the experiments, and wrote the manuscript. I.M. performed all *Drosophila* experiments and data analysis, except behavioral tests and pyruvate measurements. M.B. conducted behavioral experiments and data analysis. Z.K.A., M.D.W., and S.A. performed RNAseq analysis. C.P.C. and J.D.J.S. performed all hippocampal neuron experiments and analysis. N.A. and O.C. performed all COS7 cell experiments and analysis. P-Y.P. and T.P. performed pyruvate measurements. N.D. and S.F.T. provided technical assistance.

## STAR METHODS

### EXPERIMENTAL MODEL AND SUBJECT DETAILS

*Drosophila melanogaster* were raised on a standard cornmeal/agar diet (8g Agar, 60g cornmeal, 50g yeast, 20g glucose, 50g molasses, 19ml ethanol, 1.9g Nipagin and 10ml propionic acid in 1L of water). Animals were raised in groups up to 20 until 5 days old at 25°C in a 12/12-hour light/dark regime at 60% humidity.

For developmental depletion of *dTzap*, flies were kept at 29°C until eclosion, and adults after hatching were transferred to 18°C and kept there for 10-14 days. For the reverse experiment of adult only depletion of *dTzap* (quantifications shown in F) flies of the same genotypes were kept at 18°C until eclosion, and adults were immediately transferred to 29°C, and kept there for 10-14 days.

For the behavioral experiment on day 5, the wings were cut under CO_2_ anesthesia. They were left to recover for 48h within individual containers with access to fresh food before being transferred to the experimental set-up.

#### Cryosections of adult fly heads

Cryosections were prepared for further Laser Micro Dissection of DCN clusters. Fly heads were cut from the body and aligned in plastic molds (on ice) filled with Tissue-Tek O.C.T. Compound and immediately frozen on dry ice. Before sectioning blocks were kept at −20°C in a cryostat chamber for 10min to adjust their temperature and avoid cracking during trimming. 15µm frozen sections were performed using Leica CM3050S Cryostat. Sections were put on PEN-Membrane Slides (Leica, #11505189) and kept at −80°C in boxes containing Silica gel beads until the LMD procedure.

#### Laser Micro Dissection

Was performed from the dehydrated brain cryosections following described protocol ^35, 71^ followed by total RNA isolation. DCN clusters were visualized on cryosections under UV light by two fluorescence markers (CD8::GFP in membranes, and RedStinger in cell nucleus). Control samples were formed from random cuts of central brain regions from the same brains. Dissected tissue was collected (∼ 25 clusters/sample) into the RNAse free PCR-tubes containing 0.2% Triton X100 in DEPC treated water (Invitrogen #AM9916) with 1µl of RNases inhibitor solution (RNaseOUT Recombinant Ribonuclease Inhibitor, Quiagen #10777019). Samples were stored at −80°C until RNA isolation.

Total RNA was isolated using Ambion RNaqueous Micro Kit (Invitrogen) adapted for LMD samples. RNA concentration was checked using TapeStation, High Sensitivity D1000 Screen Tape (Agilent).

#### RNA sequencing

mRNA library preparation was realized following the manufacturer’s recommendations using SMART-Seq v4 Ultra Low Input RNA Kit from TAKARA with Nextera Tagmentation Kit. Final samples pooled library preps were sequenced on NGS NextSeq 500 Illumina using a High Output cartridge (2x400Millions of 150 bases reads). Corresponding to 2x25Millions of reads per sample after demultiplexing.

#### RNA sequencing data analysis

Reads were aligned to the *Drosophila melanogaster* genome (r6.10) using STAR (2.5.4) ^72^. We identified differentially expressed genes in adult clusters (Figure 1) by performing a Mann-Whitney U test on each DCN sample compared to the random central brain cuts samples. We used a multidimensional scaling analysis to exclude poorly clustering outlier samples. p-Values were adjusted using via Benjamini-Hochberg procedure (Supplemental table S1). Putative *ato*-targets were found using i-cisTarget software. Genes with an adjusted p-value < 0.01, base pair reads > 100bp, and 2LogFold Change > 2 were used in the follow-up analysis. The upregulated gene list was manually curated by gene function annotation obtained from FlyBase and used for RNAi-screen (Supplemental Figure S1).

RNA-seq data analysis was performed with R and RStudio (DESeq2 package). Sequencing reads and pre-processed sequencing data are available in the NCBI Gene Expression Omnibus.

#### qRT-PCR (*Drosophila* brains)

Total RNA for qRT-PCR analysis was isolated using column-based RNeasy Micro Kit (Qiagen) from dissected *Drosophila* L3 brains (10-15 brains/sample), and beads-based RNAdvance Cells v2 Kit (Beckman Coulter) from adult fly heads (10-15 heads/sample) following the manufacturer’s protocol. Reverse transcription was performed starting with 500 ng/µl total RNA concentration, by using one step Verso cDNA Synthesis Kit (Thermo Fisher Scientific). qPCR was performed on QPCR LC480 cycler (Roche), using Light Cycler 480 SYBR Green I Master Mix (Roche), and self-designed and tested primers (Key reagents table). The experiments were performed with at least three independent biological samples and carried out with technical triplicates. Data are shown as fold change, calculated as 2^-DDCt^ using *Act88F* as a calibrator gene.

#### Immunofluorescence of *Drosophila* brains

For brain immunofluorescence labeling L3 instant larvae brains and 3-7 days old fly brains were dissected in 1X-PBS and transferred to a tube for fixation in 4% paraformaldehyde in PBS for 20 min at room temperature. Brains were washed in PBS (one time) and 0.3% Triton-X (PBST) three times for 10 min each. Next, brains were placed in blocking solution (2% normal bovine serum (Sigma-Aldrich) in PBST) overnight at 4 °C. The samples were then incubated in primary antibodies diluted in working solution (1 ml of 2% BSA, 250µl 10% NatAzid in 50ml 2% PBST) overnight at 4 °C. Used primary and secondary antibodies are listed in the Key Reagents table. Brains were then washed three times in PBST for 10 min each and incubated with a fluorochrome-conjugated secondary antibody overnight at 4 °C (listed in the Key Reagents table). The brains were washed three times in PBST for 10 min each. Finally, the samples were mounted onto microscope slides with Vectashield Vibrance Antifade Mounting Medium (Vector Laboratories) for subsequent analysis using the SP8 Leica or Olympus FV1200 confocal microscopes. Images were analyzed using the Fiji ImageJ and IMARIS software.

Trans-TANGO experiment was performed with the DCN-specific *ato-Gal4^14a^* driver line, and the number of postsynaptic neurons was counted manually from their cell bodies, including all cells with weak or strong labeling to reveal all potential connections.

pHluorin to mCherry fluorescence ratio detection was made in freshly dissected brains in PBS on ice. After dissection brains were immediately mounted on the Superfrost slides with Vectashield Antifade Mounting Medium (Vector Laboratories) and imaged on Confocal SP8 Leica.

#### Behavioral arena

The behavioral arena used is a modification of Buridan’s Paradigm ^55^. The arena consists of a round platform of 119 mm in diameter, surrounded by a water-filled moat (Figure 6H). The arena was placed into a uniformly illuminated white cylinder. The setup was illuminated with four circular fluorescent tubes (Philips, L 40w, 640C circular cool white) powered by an Osram Quicktronic QT-M 1×26–42. The four fluorescent tubes were located outside of a cylindrical diffuser (Canson, Translucent paper 180gr/m2) positioned 145 mm from the arena center. The temperature on the platform during the experiment was maintained at 25°C.

#### ChiPseq available data analysis

Available ChiPseq data from whole-organism adult *Drosophila* females were available at https://www.encodeproject.org/experiments/ENCSR472BOK/ ^41^. Human ChiPseq data were published by *Jahn et al., 2017* ^42^. Drosophila reads were aligned with the dm6 reference *Drosophila* genome, human – with hg38 reference genome using Integrative Genome Viewer IGV and Genomatix software.

#### Pyruvate accumulation measurements

Experiments were done as described by Plaçais et al., 2017 ^73^. UAS-Pyronic sensor was expressed in DCNs using the *ato-Gal4^14a^* driver.

#### Statistical analysis

Data from behavioral experiments were analyzed using R ^74^, and from other experiments – using Prism-GraphPad software. In behavior analysis transition plots were done as described before ^55^. Briefly, the platform was divided into 60*60 hexagons and the fly’s position raised the count of each hexagon by one in the arena. The scale starts at 0 (blue) and goes up till a value is calculated by the 95%-quantile of the count distribution (red). For the statistical analyses, we first checked for normal data distribution using the Shapiro-Wilk normality test. Then, we chose the appropriate parametric or non-parametric test. In the analysis of behavioral data, we used Ordinary one-way ANOVA with the Kruskal-Wallis test. In other experiments, we used Ordinary one-way ANOVA with Dunnett and Bonferroni corrections for multiple comparisons and unpaired Students t-test with Welch’s correction. *\* p<0.05, ** p*≤*0.01, *** p*≤*0.001, **** p*≤*0.0001, ns* – not significant.

### Studies in mammalian cells

#### Cell culture and transfection

COS7 cells (ATCC) were grown in Dulbecco’s modified Eagle medium (DMEM, Gibco) supplemented with 10% fetal bovine serum (FBS), 1% L-glutamine (Invitrogen), and 1% penicillin-streptomycin (Invitrogen) and tested monthly for mycoplasma contamination (MycoAlert Mycoplasma Detection Kit, Lonza). Cells were plated in 8-well glass bottom Ibidi (80827; Ibidi), at a density of 1.5 × 10^4^ per well precoated with poly-L-lysine (1 mg/L, Sigma). The following day, cells were transfected with 0.1 µg/well of pcDNA3-HA, 0.1 µg/well of pCB6-HA-mitoGFP ^75^, and 10 pmol/well of TZAP stealth siRNA (s6568, ThermoFisher) or AllStars negative control (Qiagen 1027281), with Lipofectamine 2000 (0.5 µl/well, Invitrogen) in 50 µl of Opti-MEM I (Gibco), without medium change, according to the manufacturer’s instructions.

#### Microscopy and image analysis

After 48 hours, Z-stack projection images were captured in life-imaging with a Spinning disk CSU-X1 confocal microscope (Leica) driven by MetaMorph software (63X immersion oil objective, N.A. 1.4), and processed with ImageJ software. Cells expressing MitoGFP in images were manually categorized according to mitochondrial network morphology, classified as fissioned; donut-like; fusioned and mixed (with no predominant type of morphology). The results are expressed as mean values ± SEM from three independent experiments.

For the quantitative analysis of the morphology of the mitoGFP-positive mitochondrial objects, a manual threshold mask was applied on each of 25-30 images per condition, using ImageJ software (NIH, Bethesda, Maryland). For each mitochondrial object, area, perimeter, aspect ratio, and form factor were calculated. Aspect ratio is the ratio between the major and minor axes of each mitochondrial object, and represents its length; form factor is calculated as perimeter2/(4π × area) and thus represents a combined evaluation of the length and degree of branching of the mitochondrial network. We analyzed a total of 940-1200 objects per condition from three independent experiments. The results are expressed as means ± SEM.

#### Quantitative analysis of transcript levels

To evaluate the efficiency of the siRNA-mediated silencing approach for endogenous TZAP and expression levels of genes involved in the regulation of mitochondrial dynamics, total RNA was extracted from COS7 cells, using RNeasy Micro kit (Qiagen), according to the manufacturer’s instructions, and treated with DNAse I (Qiagen) for 20 min at room temperature. RNA concentrations were determined using Nanodrop 2000c (THERMO Scientific). Complementary DNA was generated from 500 ng RNA with random hexamers and Superscript II reverse transcriptase (THERMO Fisher Scientific). Real-time PCR was performed with the LightCycler 480 qPCR system (Roche) with SYBR green detection and each of the following primer pairs in 5′-to 3′ orientation (see Key reagents table). Relative gene expression levels were calculated using the 2−ΔΔCt method, with GAPDH used as the reference gene for normalization. Statistical analyses were performed using GraphPad Prism 7 software. Normal distribution was checked using the D’Agostino Pearson normality test before statistical comparisons. Data were analyzed by two-way ANOVA. When pertinent, the Tukey post hoc test was used for multiple comparisons. The sample size was standard and determined based on previous experience. Analyses were not blinded and no sample was excluded. The results are presented as means ± SEM from three independent experiments.

### Studies in rat primary hippocampal neurons

#### Animals

Wild-type rats were of the Sprague-Dawley strain Crl: CD (SD) and were bred by Janvier Labs (France) following the international genetic standard protocol (IGS). All procedures relating to the care and treatment of animals were performed following the guidelines of the European Directive 2010/63/EU and the French Decree n° 2013-118 concerning the protection of animals used for scientific purposes.

#### Primary co-culture of postnatal hippocampal neurons and astrocytes

Imaging experiments for mitochondrial Ca^2+^ dynamics and glutamate release were performed in primary co-cultures of hippocampal neurons and astrocytes using P0 to P2 rats of mixed gender. After dissection to isolate the hippocampus, cells were plated on cloning cylinders of 4.7 mm diameter attached to poly-ornithine-coated coverslips (38,000-40,000 cells per cylinder), transfected 7 days after plating using calcium phosphate as previously described^76^. Neurons were maintained in culture media composed of MEM (Thermo Fisher Scientific, #51200087), 20 mM Glucose (Sigma, #G8270), 0.1 mg/mL transferrin (Sigma, #616420), 1% GlutaMAX (Thermo Fisher Scientific, #35050061), 24 μg/mL insulin (Sigma, #I6634), 10% FBS (Thermo Fisher Scientific, #10082147) and 2% N-21 (Bio-techne, #AR008). After 5 days in vitro, neurons were incubated with media of the same composition but using only 5% FBS and adding 2 μM cytosine β-d-arabinofuranoside to curb glial growth (Sigma, #C6645). Cultures were incubated at 37°C in a 95% air/5% CO_2_ humidified incubator for 14– 21 days prior to use for imaging.

#### Plasmid Constructs for live cell imaging in rodent neurons

The following DNA constructs were used for expressing biosensors for live cell imaging: mito^4x^-GCaMP6f ^47^ and iGluSnFR3, which was a gift of Kaspar Podorgski (Addgene plasmid #178330) ^77^. The shRNA sequence for the Zbtb48 gene (ENSRNOG00000009595.6) was generated from GPP Web Portal (https://portals.broadinstitute.org/gpp/public/seq/search). The selected shRNA sequence 5-’AGGGATGGTGATGGCGATTAT-3’ was cloned into pLKO.1-TRCmTagBFP2, a gift from Timothy A. Ryan (Addgene plasmid #191566), through AgeI and EcoRI sites as previously described (Moffat et al Cell. 2006). The primers to generate the insert containing the shRNA sequence are annotated in the Key Resources table. For control experiments was used the same pLKO-mTagBFP2 plasmid without the shRNA insert.

#### Live imaging of neurons

Neuronal imaging was performed in continuously flowing Tyrode’s buffer containing (in mM) 119 NaCl, 2.5 KCl, 1.2 CaCl_2_, 2.8 MgCl_2_, 20 glucose, 10 µM 6-cyano-7-nitroquinoxaline-2,3-dione (CNQX) and 50 µM D,L-2-amino-5-phospho-novaleric acid (AP5), buffered to pH 7.4 at 37°C using 25 mM HEPES. Imaging was performed on a custom-built laser-illuminated epifluorescence microscope (Zeiss Axio Observer 3) coupled to an Andor iXon Ultra camera (model #DU-897U-CSO-#BV), whose chip temperature is cooled down to −90 °C using the Oasis™ UC160 Cooling System allowing the reduction of noise in the measurements. Illumination using a fiber laser of wavelength 488 (Coherent OBIS 488nm LX 30mW) was controlled by a custom Arduino-based circuit coupling imaging and illumination. Primary hippocampal neurons were grown in coverslips (D=0.17mm, Warner Instruments, #640705), mounted on an imaging chamber for field stimulation (Warner Instruments, #RC-21BRFS), and imaged through a 40x Zeiss oil objective “Plan-Neofluar” with an NA of 1.30 (WD=0.21mm). For glutamate release measurements using iGluSnFR3 data was obtained from imaging at 350Hz during single action potentials induced by 1ms field stimulation. In the case of mitochondrial Ca^2+^ assays using mito^4x^-GCaMP6f, data was obtained from imaging at 5 Hz. Mito^4x^-GCaMP6f may mislocalize occasionally in the cytosol. To avoid inaccuracies in peak response detection, we extracted mitochondrial Ca2+ peak responses after cytosolic Ca2+ clearance, as previously described ^47^. The temperature of all experiments was clamped at 37 °C and was kept constant by heating the stimulation chamber through a heated platform (Warner instruments, #PH-2) together with the use of an in-line solution heater (Warner instruments, #SHM-6), through which solutions flowed at 0.35ml/min. The temperature was monitored constantly using a feedback-look temperature controller (Warner instruments, #TC-344C).

#### Image analysis, statistics, and exclusion criteria

Images were analyzed using the ImageJ plugin Time Series Analyzer where ∼15-20 (for mitochondrial *Ca^2+^) or* ∼3-4 (for glutamate release) regions of interest (ROIs) per neuron were selected and the fluorescence was measured over time. Statistical analysis was performed with GraphPad Prism v8.0 for Windows using the nonparametric Mann-Whitney U test to determine the significance of the difference between two unpaired conditions, as we did not assume that the distributions of our two unpaired datasets follow a normal distribution. *p < 0.05* was considered significant and denoted with a single asterisk, whereas *p < 0.01, p < 0.001,* and *p < 0.0001* are denoted with two, three, and four asterisks, respectively. The n value represents the number of cells imaged and statistic tests performed in each experiment are indicated in the respective figure legends.

Glutamate release and mitochondrial Ca^2+^ in response to electrical activity (ΔF) were normalized to the resting fluorescence (F_0_). A criteria of inclusion/exclusion data was established to avoid overestimating ΔF/F_0_ in responding neurons with low F_0_ values, setting an arbitrary threshold such that F_0_/background > 1.25 to be included for further analysis (11/49 Mito^4x^-GCaMP6f and 0/25 iGluSnFR3 were excluded). Experiments were replicated with neurons from at least 4 independent primary cultures per condition.

#### Mitochondrial shape parameters

Different morphological aspects of mitochondria were analyzed from the images taken in the mitochondrial calcium measurement experiments. Images were analyzed using the ImageJ plugin Trainable Weka Segmentation following the criteria described previously ^78^. For cultured neurons, in addition to the global background of the image not containing any cell type, pixels containing astrocytes were also selected as a secondary background, which increased specificity in the mitochondrial shaping measurements performed by the Trainable Weka Segmentation plugin. The morphological aspects of mitochondria analyzed for neurons were area, perimeter, circularity, and aspect ratio, the values were obtained in pixels and later converted to micrometers following the characteristics of the camera used (1 pixel = 0.4 µm).

## Key resources table

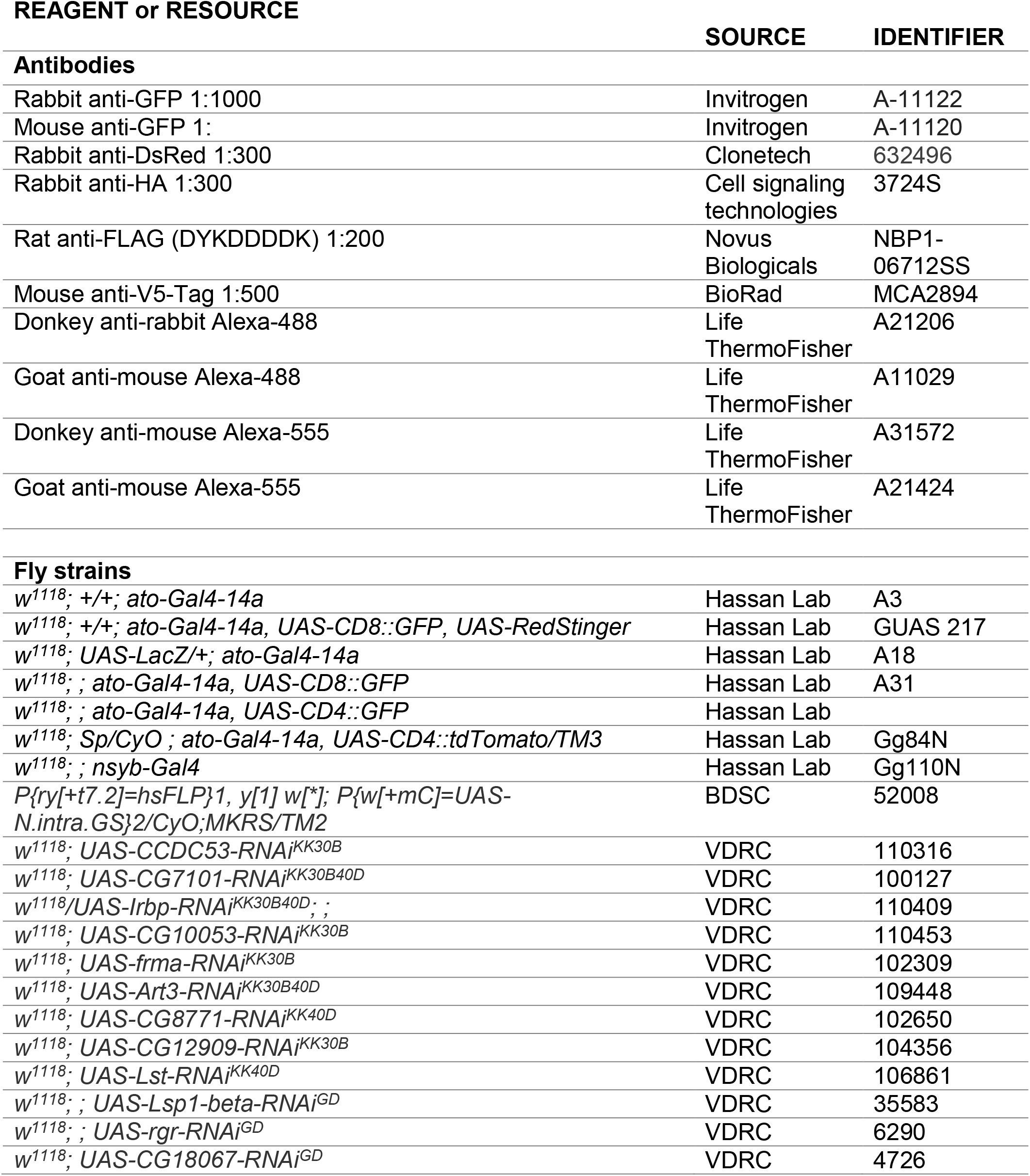
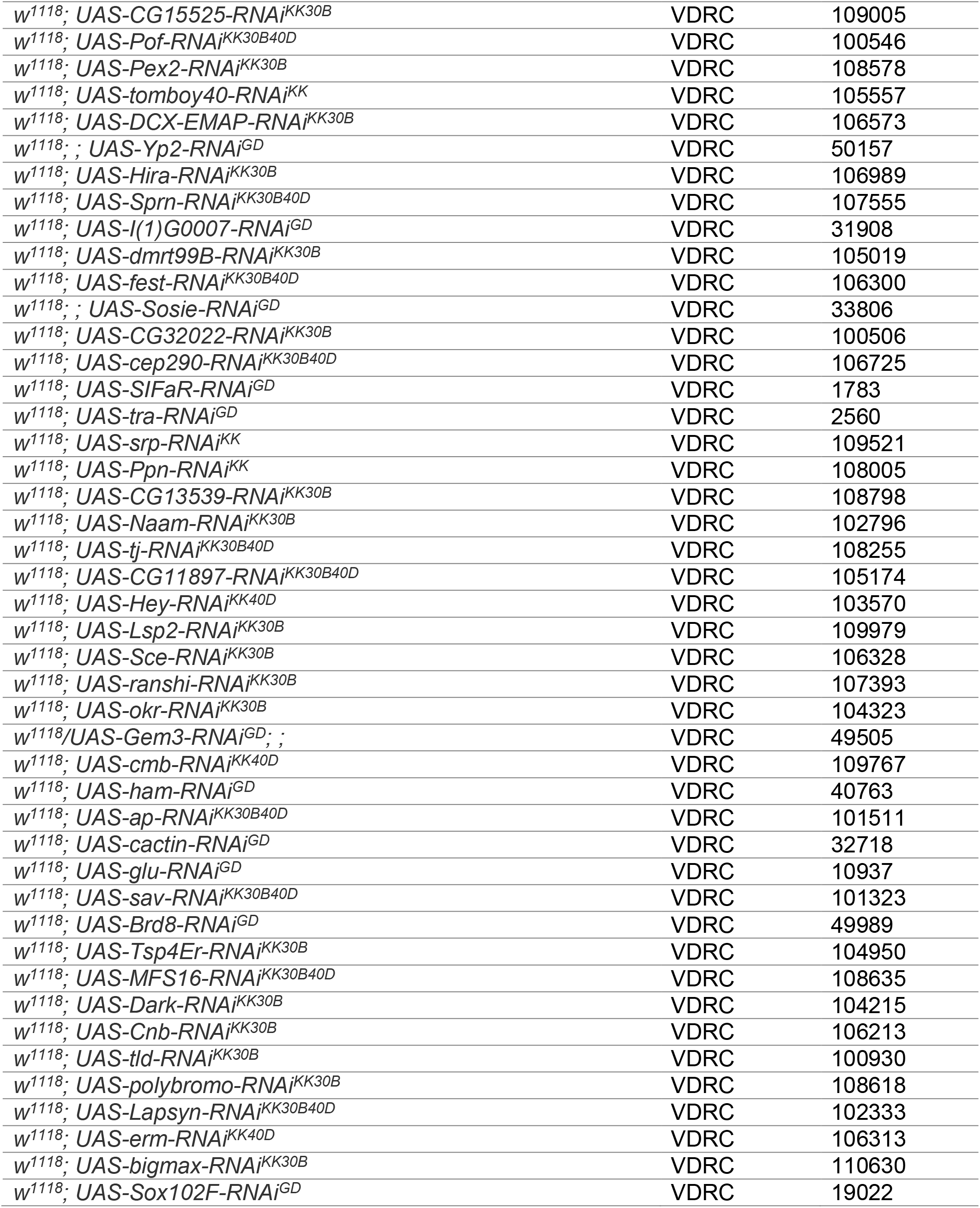
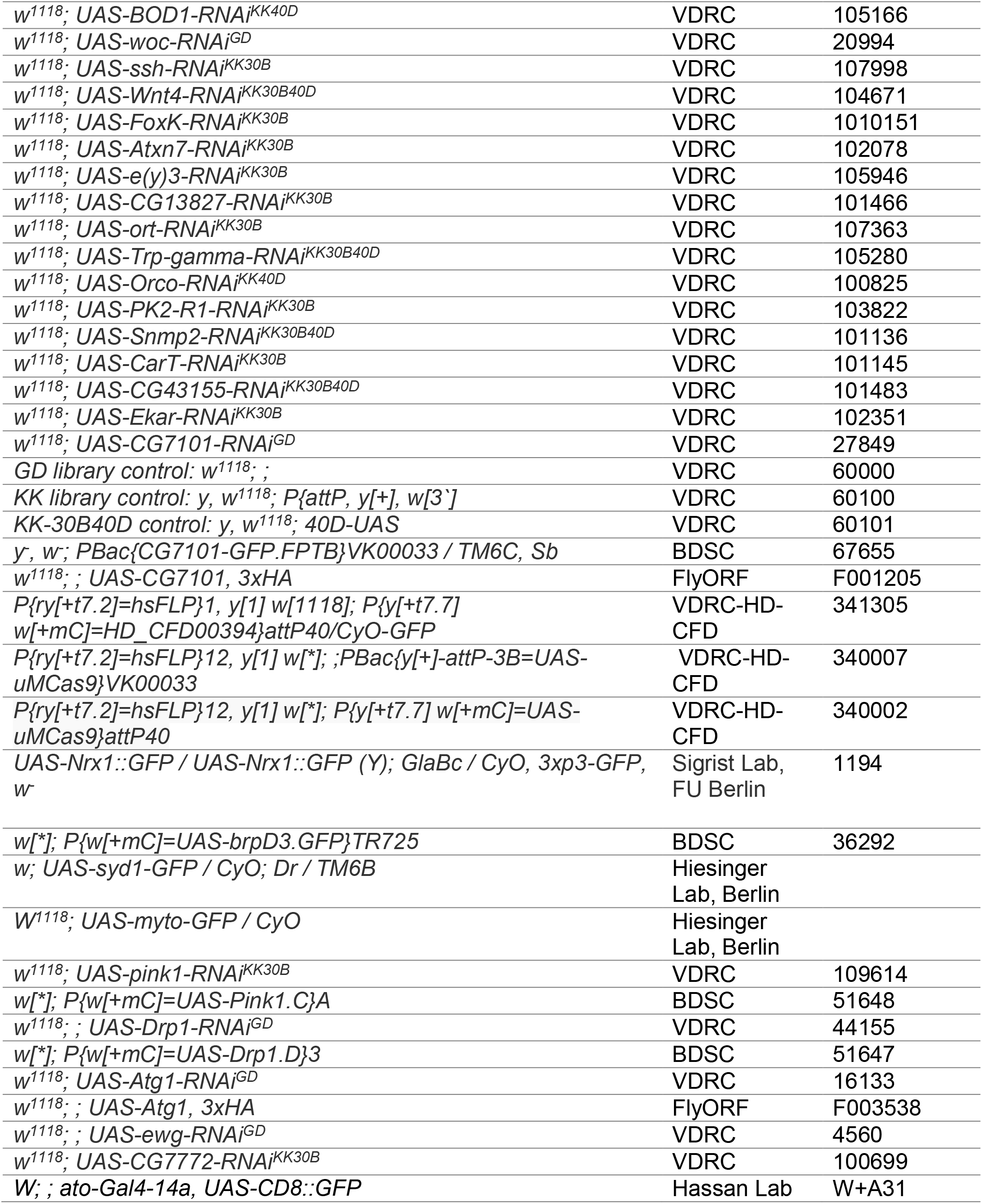
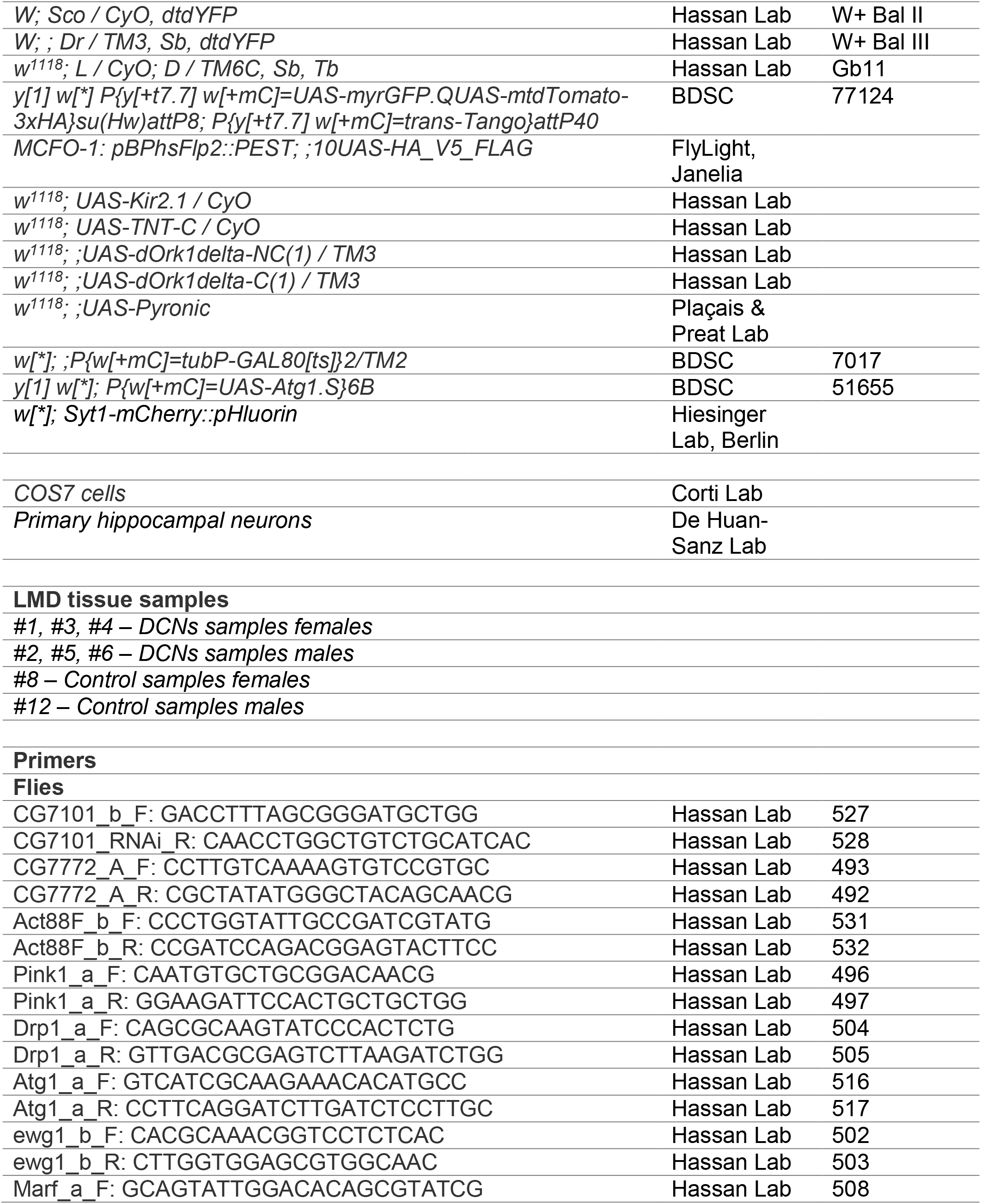
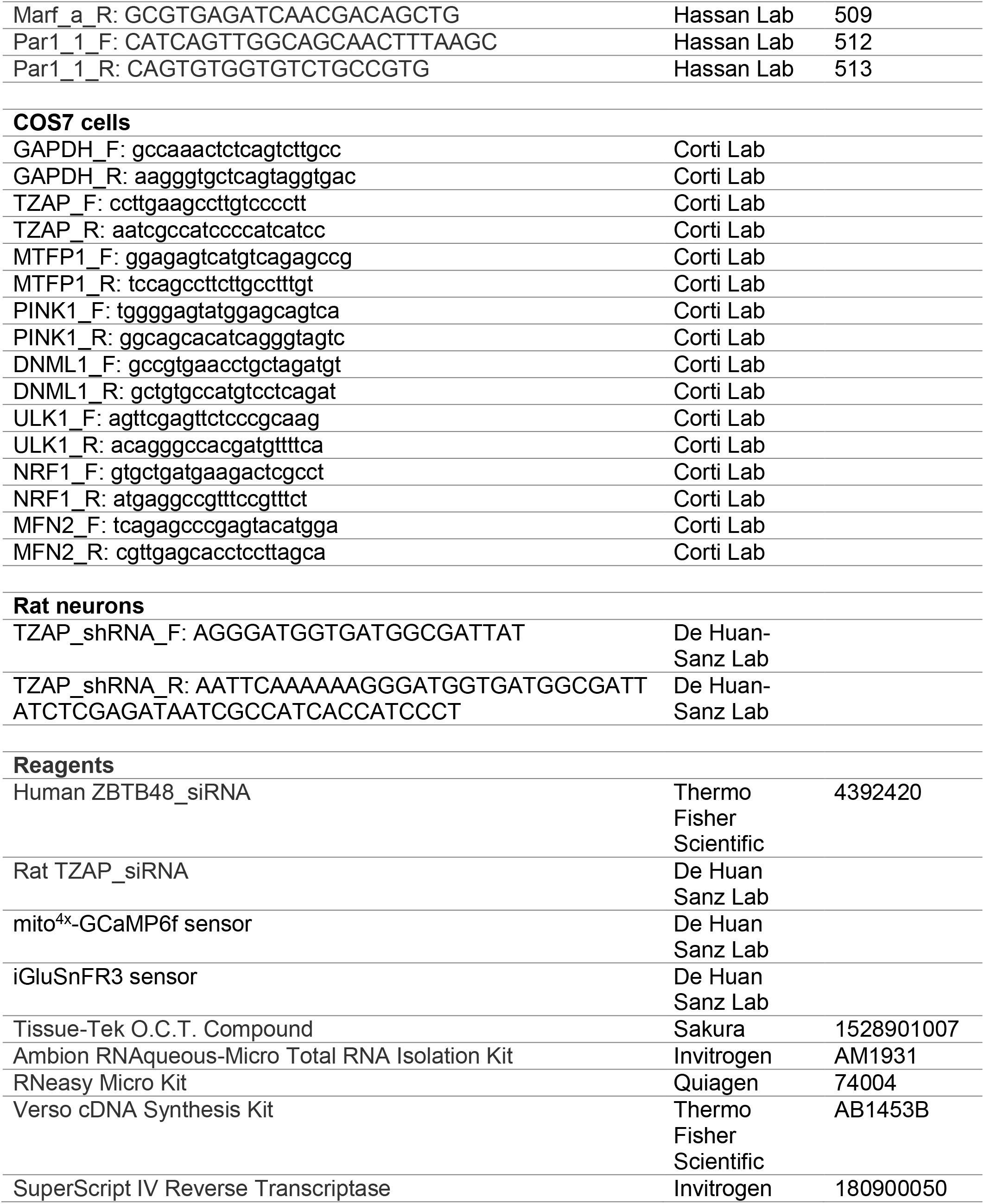
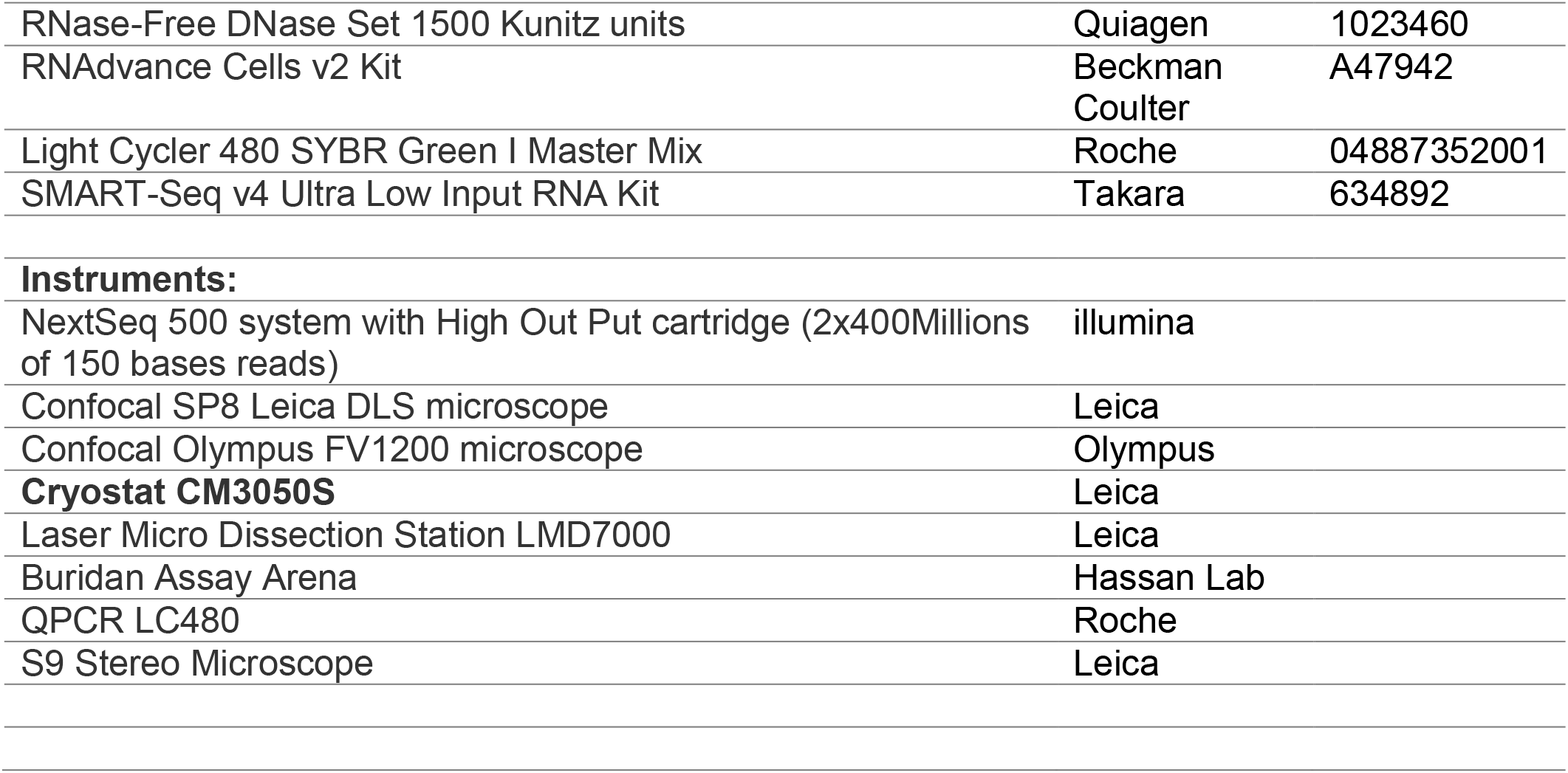

## Supplementary data file

### Supplementary File 1

Analysis of LMD dissected cell clusters (tab1), genes expressed in DCNs (tab2) and list of genes for the RNAi-screen (tab3).

**Supplementary Figure S1.**
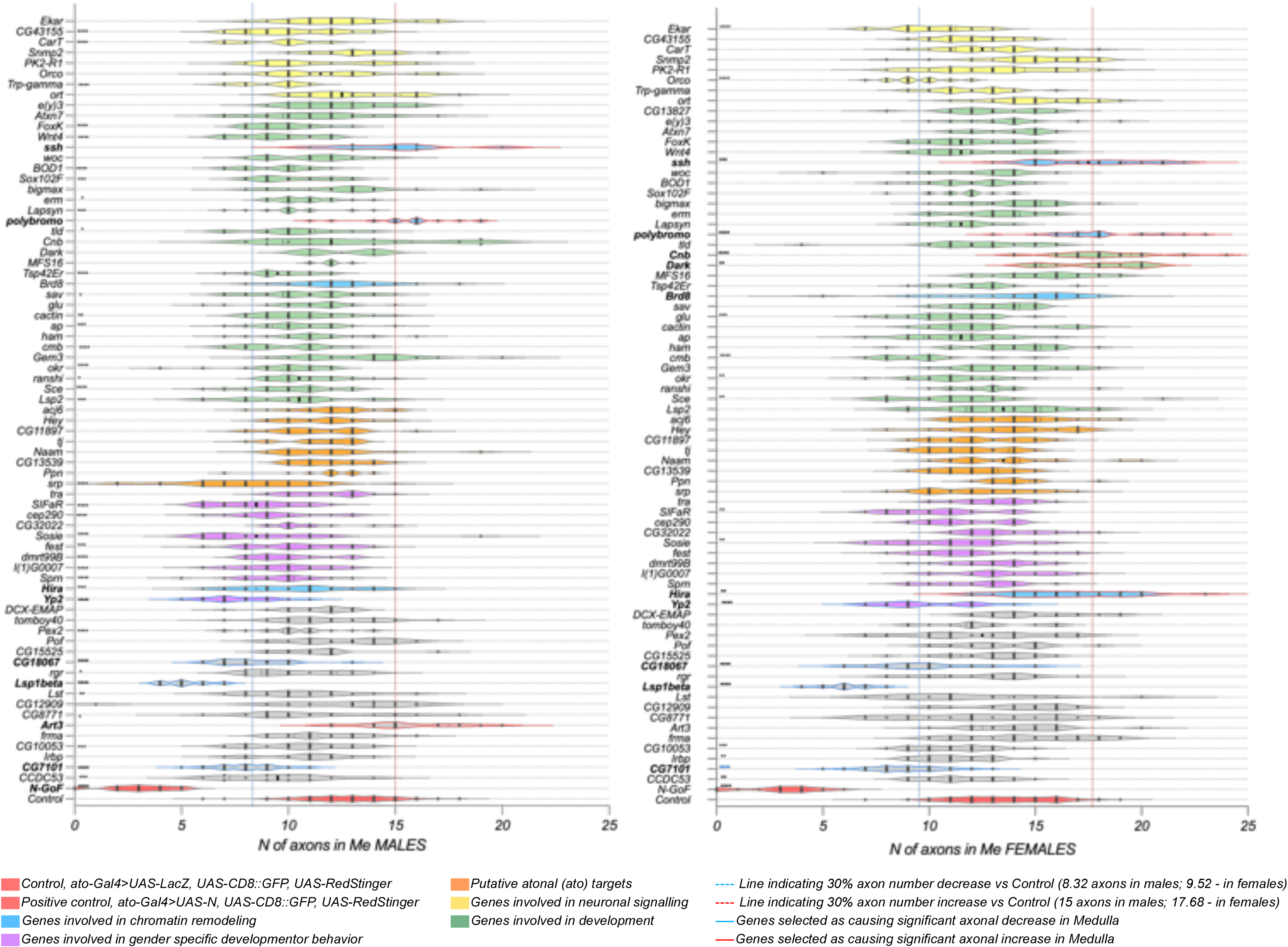
RNAi screen of 74 genes expressed in DCNs. As a readout for the screen, the number of DCNs axons in the medulla was used and compared to Control flies (*w^1118^;UAS-LacZ/+;ato-Gal4^14A^,UAS-CD8::GFP,UAS-RedStinger*). Flies with *Notch* overexpression were used as positive Control, (*N-GoF*) as it was shown earlier that Notch overexpression prevents medulla innervation. We considered observed phenotypes as positive if changes in medulla innervation were equal or more than 30% from the mean axon number in Control flies (dashed blue line corresponds to a 30% decrease and dashed red line to 30% increase values). Candidate genes (in bold) selected during the screen have blue (for decreased innervation) and red (for increased innervation) line borders on the plot. We divided genes by groups, according to known annotations of their function (corresponding colors with description). Crosses were set in groups of five RNAi lines plus Control with each group. Adult flies were dissected at 3-7 days old. The final number of analyzed optic lobes in Control flies, 158 males & 154 females. In tested genes in the screen, *n* value varies from 8 to 38 in males and from 12 to 45 in females (depending on genotype and the number of repeats). Statistical analysis was done using *Kruscal-Wallis One-way ANOVA* with Dunn’s corrections for multiple comparisons. *\* p<0.05, ** p*≤*0.01, *** p*≤*0.001, **** p*≤*0.0001, ns* – not significant.

**Supplementary Figure S2.**
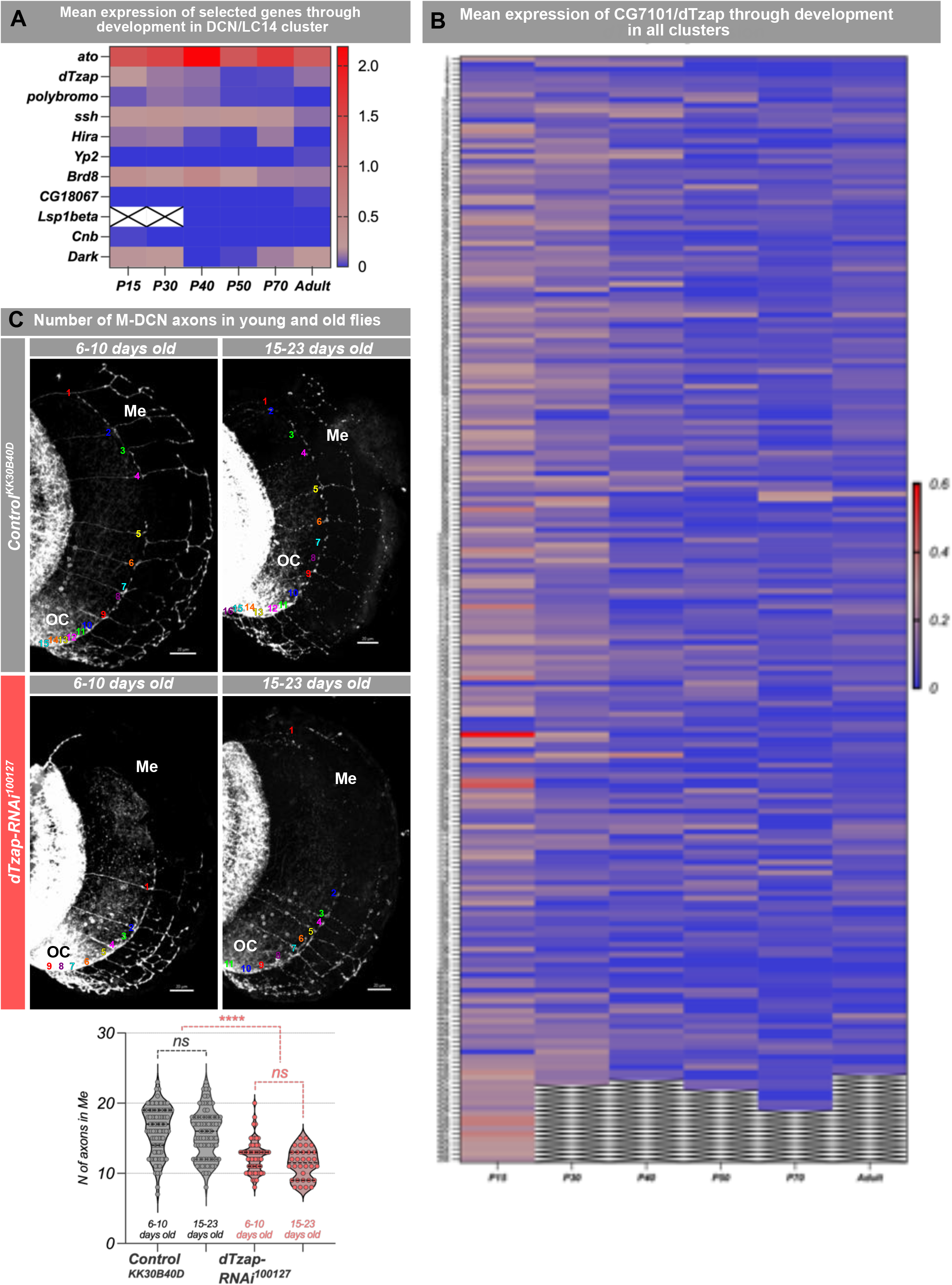
scRNAseq and phenotype of CG7101 knockdown. Heat maps were generated from available dataset of single-cell RNAseq of the fly visual system at different developmental stages. **A** – normalized mean expression of 10 candidate genes, with a significant change in the number of axons reaching medulla, in the DCN cluster at different developmental stages. **B** – normalized mean expression of *dTzap* throughout development in different neuronal clusters. **C** – Number of M-DCNs axons in young (6-10 days old) and old (15-23 days old) flies. OC – optic chiasm, Me – medulla, scale bar – 20µm. Statistical analysis was performed with GraphPad Prism 8 using the nonparametric Mann-Whitney U test. *\* p<0.05, ** p*≤*0.01, *** p*≤*0.001, **** p*≤*0.0001, ns* – not significant.

**Supplementary Figure S3.**
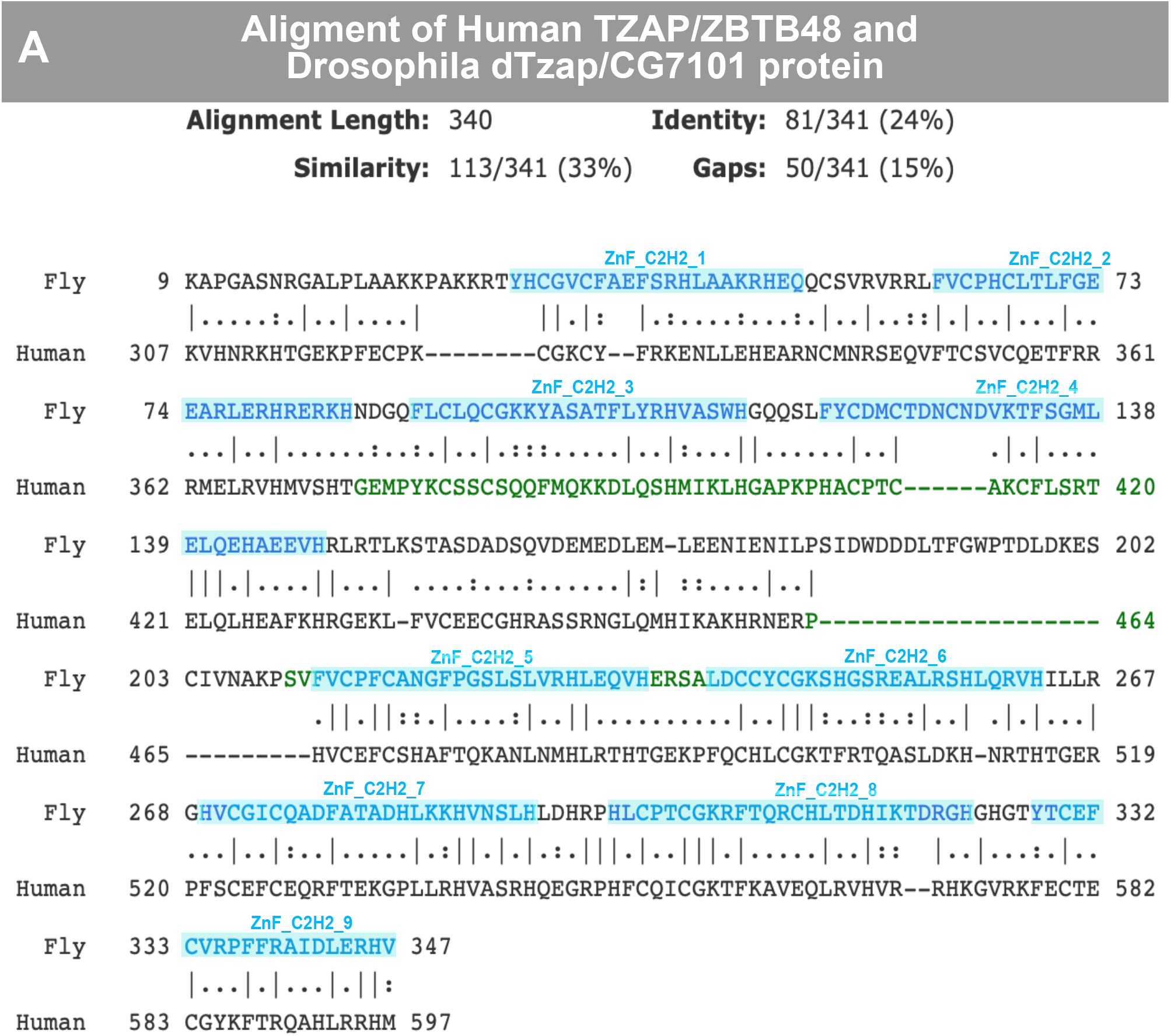
Alignment of human TZAP/ZBTB48 and *Drosophila* CG7101/dTzap protein. **A**– Alignment of human TZAP/ZBTB48 and *Drosophila* CG7101/dTzap protein shows a high level of homology. ZnF_C2H2 domains of fly protein shown in blue (alignment adopted from DIOPT, DRSC integrative ortholog prediction tool).

**Supplementary Figure S4.**
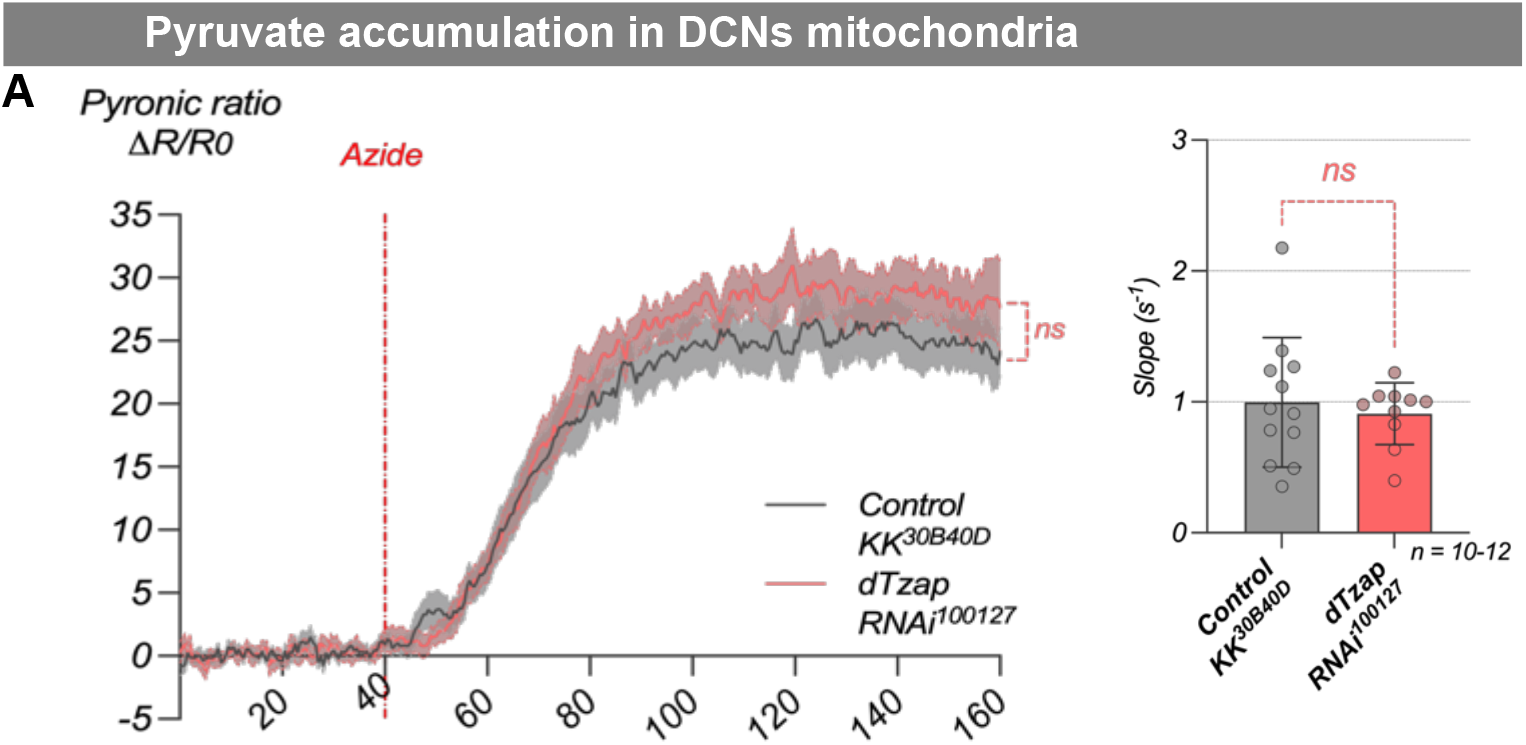
In vivo pyruvate imaging in DCNs reveals no impact of dTzap depletion on mitochondria pyruvate consumption. **A *–*** Pyruvate accumulation in DCNs triggered by sodium azide treatment in flies expressing Pyronic sensor in DCNs (*w^1118^; dTzap-RNAi^100127^; UAS-Pyronic/ato-Gal4*^14a^ vs *w^1118^; Control-KK^30b40D^; UAS-Pyronic/ato-Gal4^14a^*). A’ – Calculations of pyruvate accumulation slope (dynamic) and plateau. Statistical analysis was performed with GraphPad Prism 8. Normal distribution was checked using the D’Agostino Pearson normality test before statistical comparisons. Unpaired T-test with Welch’s correction was used *\* p<0.05, ** p*≤*0.01, *** p*≤*0.001, **** p*≤*0.0001, ns* – not significant.

**Supplementary Figure S5.**
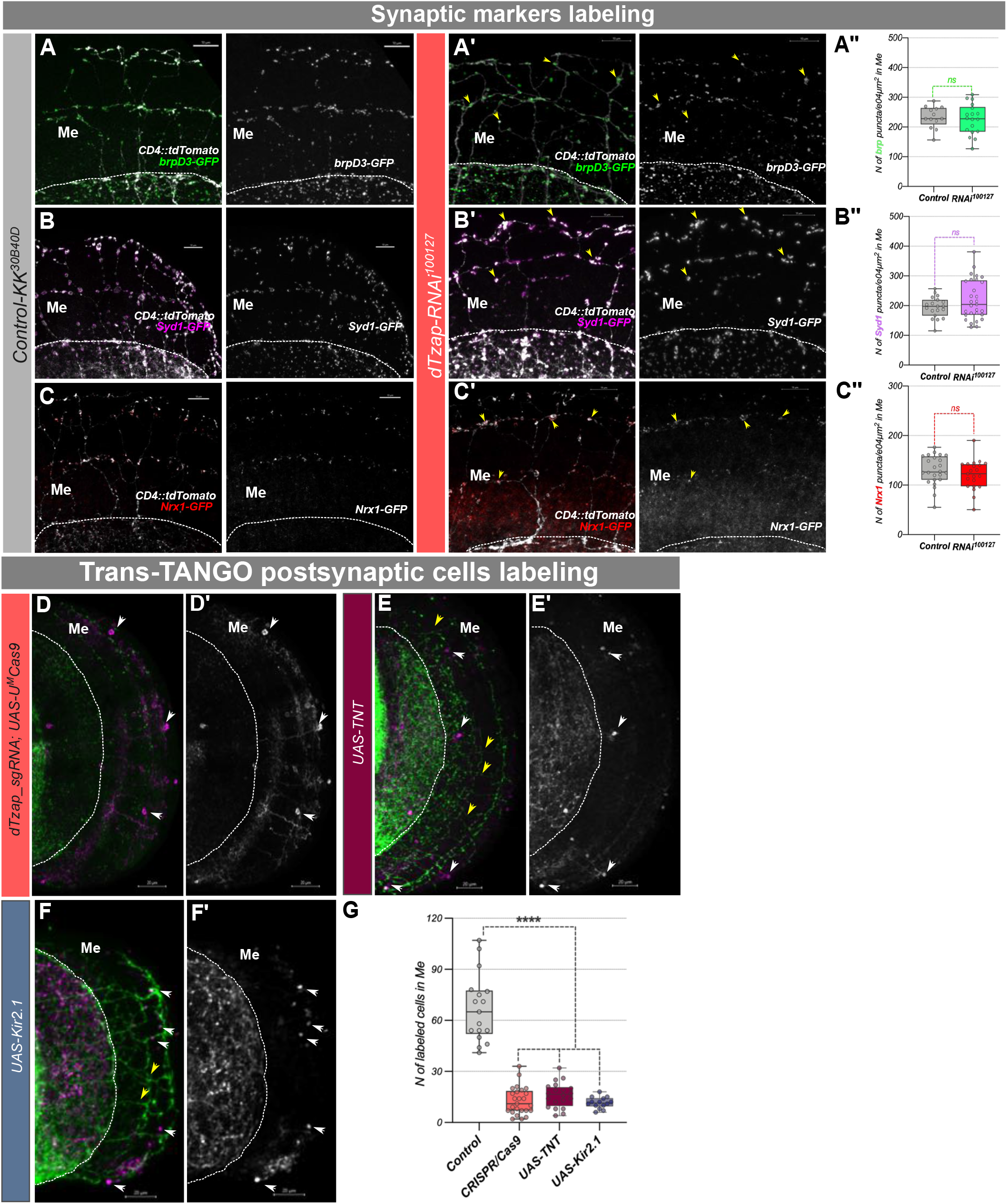
Presynaptic marker localization does not require dTzap, and TransTango is activity dependent. **A-C’’**– Labeling of presynaptic markers in M-DCNs axons in *dTzap-RNAi^100127^* background showed no significant difference to controls (*w^1118^; UAS-KK^40D30B^*), scale bar 10µm. Three major synaptic markers (brpD3, Syd1, and Nrx1) were present at the presynaptic sites in quantities comparable to control condition (A”, B”, C”). GFP positive puncta in the medulla were automatically counted using IMARIS software, and normalized to corresponding medulla area. Puncta counting: BrpD3 – 13 optic lobes (control) and 18 optic lobes (*RNAi^100127^*), Syd1 – n=19 optic lobes (control) and 29 optic lobes (*RNAi^100127^*), Nrx1 – n=24 optic lobes in (control) and 21(*RNAi^100127^*). **D-F’ *–*** TransTango anterograde postsynaptic labeling of DCNs connectivity in case of *dTzap* downregulation using CRISPR/Cas9 approach (*UAS-myrGFP.QUAS-mtdTomato-3xHA;transTANGO/UAS-dTzap_sgRNA;ato-Gal4^14a^/UAS-U^M^Cas9^340007^)* (A-A’’), and DCNs silencing with Tetanus toxin (B, B’, n=20 optic lobes) and potassium channel overexpression with *UAS-Kir2.1* (F, F’, n=14 optic lobes). DCNs expressing *myrGFP* on the cellular membrane are shown in green (D, E, F) and postsynaptic labeled cells in magenta (merged picture) and grayscale (D’, E’, F’). Postsynaptic connectivity was analyzed by manually counting cell bodies in medulla (white arrowheads) labeled with dsRed fluorescence (magenta, merged picture – D, E, F, and grayscale – D’, E’, F’), from their cell bodies, including all cells with weak or strong labeling to reveal all potential connections. Although there are still M-DCNs axons visible in medulla (yellow arrowheads) they fail to establish connectivity with neighboring cells. Scale bar 20µm. Statistical analysis was done using an unpaired Student’s t-test with Welch’s correction for axon counting, and one-way ANOVA with Bonferroni corrections for multiple comparisons in TransTango labeling analysis. *\* p<0.05, ** p*≤*0.01, *** p*≤*0.001, **** p*≤*0.0001, ns* – not significant.

**Supplementary Figure S6.**
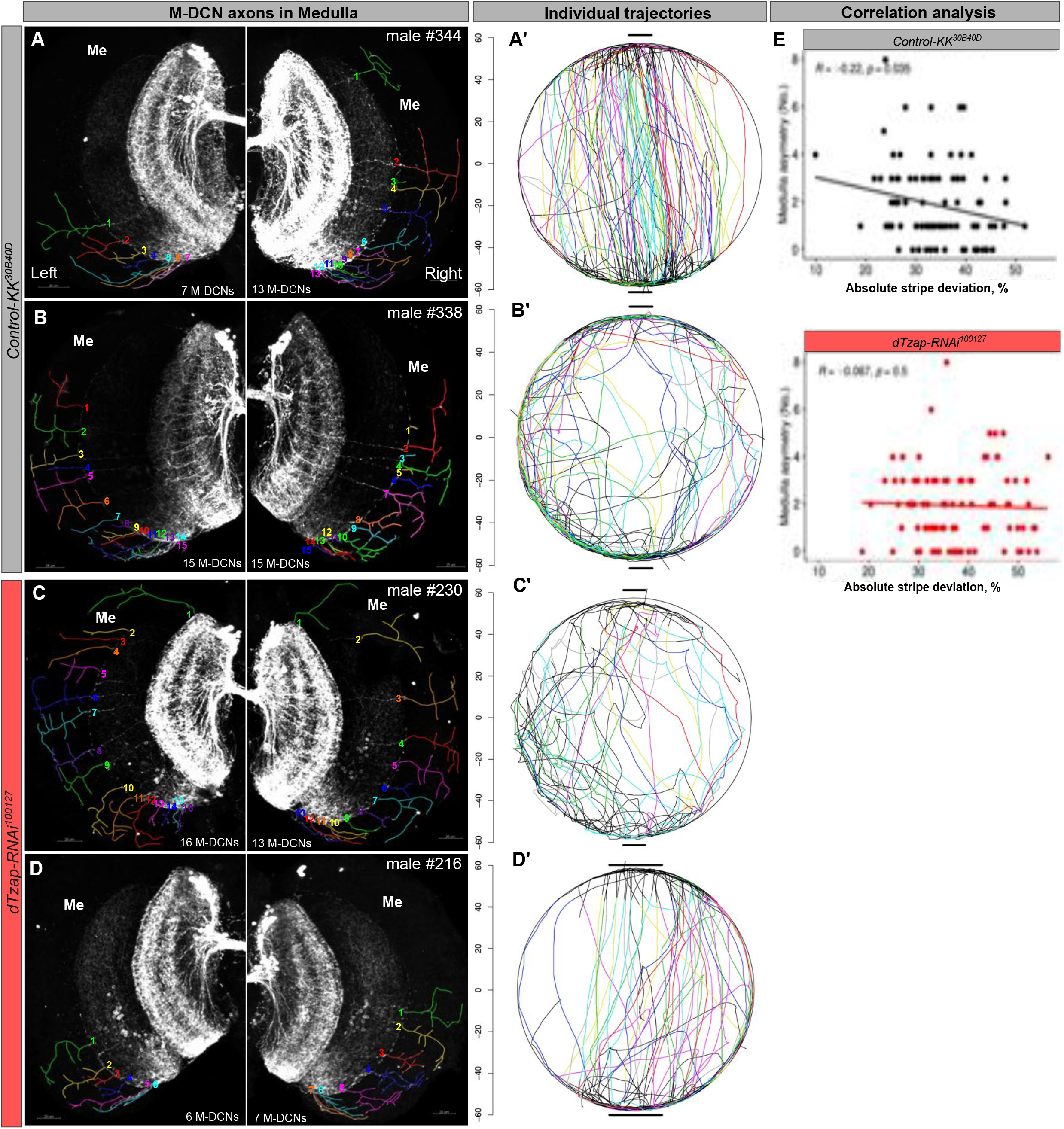
Correlation analysis of Individual trajectories in Buridan assay and DCN circuit anatomy. **A, D**– DCN neurons in individual control males (#344 and #338, *W+;UAS-KK^40D30B^; ato-Gal4^14A^,UAS-CD4::GFP)* and males with *dTzap* downregulation (#230 and #216, *W+;UAS-dTzap-RNAi^100127^;ato-Gal4^14A^,UAS-CD4::GFP)* showing the different degrees of asymmetry. **A’-D’** – Corresponding individual behavioral paths in Buridan assay. **E** – Correlation analysis of individual DCN innervation asymmetry in medulla and absolute stripe deviation (125 control flies and 135 *RNAi^100127^* flies. Both, males and females in equal ratio). Me – medulla, Lo – lobula, OC – optic chiasm. Statistical analysis was done using one-way ANOVA with the Kruskal-Wallis test for the behavioral experiment, correlation analysis was performed using Spearman rank correlation.

## REFERENCES

1. Hassan, B.A., and Hiesinger, P.R. (2015). Beyond Molecular Codes: Simple Rules to Wire Complex Brains. Cell 163, 285–291. 10.1016/j.cell.2015.09.031.

2. Sanes, J.R., and Zipursky, S.L. (2020). Synaptic Specificity, Recognition Molecules, and Assembly of Neural Circuits. Cell 181, 536–556. 10.1016/J.CELL.2020.04.008.

3. Isshiki, T., Pearson, B., Holbrook, S., and Doe, C.Q. (2001). Drosophila Neuroblasts Sequentially Express Transcription Factors which Specify the Temporal Identity of Their Neuronal Progeny. Cell 106, 511–521. 10.1016/S0092-8674(01)00465-2.

4. Pichaud, F., Treisman, J., and Desplan, C. (2001). Reinventing a common strategy for patterning the eye. Cell 105, 9–12. 10.1016/S0092-8674(01)00292-6.

5. Sagner, A., Zhang, I., Watson, T., Lazaro, J., Melchionda, M., and Briscoe, J. (2021). A shared transcriptional code orchestrates temporal patterning of the central nervous system. PLoS Biol 19, e3001450. 10.1371/JOURNAL.PBIO.3001450.

6. Xie, Q., Li, J., Li, H., Udeshi, N.D., Svinkina, T., Orlin, D., Kohani, S., Guajardo, R., Mani, D.R., Xu, C., et al. (2022). Transcription factor Acj6 controls dendrite targeting via a combinatorial cell-surface code. Neuron 110, 2299–2314.e8. 10.1016/J.NEURON.2022.04.026.

7. Nern, A., Zhu, Y., and Zipursky, S.L. (2008). Local N-Cadherin Interactions Mediate Distinct Steps in the Targeting of Lamina Neurons. Neuron 58, 34–41. 10.1016/j.neuron.2008.03.022.

8. Langen, M., Agi, E., Altschuler, D.J., Wu, L.F., Altschuler, S.J., and Hiesinger, P.R. (2015). The Developmental Rules of Neural Superposition in Drosophila. Cell 162, 120–133. 10.1016/J.CELL.2015.05.055.

9. Langen, M., Koch, M., Yan, J., De Geest, N., Erfurth, M.L., Pfeiffer, B.D., Schmucker, D., Moreau, Y., and Hassan, B.A. (2013). Mutual inhibition among postmitotic neurons regulates robustness of brain wiring in Drosophila. Elife 2. 10.7554/ELIFE.00337.

10. Sales, E.C., Heckman, E.L., Warren, T.L., and Doe, C.Q. (2019). Regulation of subcellular dendritic synapse specificity by axon guidance cues. Elife 8. 10.7554/ELIFE.43478.

11. Özel, M.N., Kulkarni, A., Hasan, A., Brummer, J., Moldenhauer, M., Daumann, I.M., Wolfenberg, H., Dercksen, V.J., Kiral, F.R., Weiser, M., et al. (2019). Serial Synapse Formation through Filopodial Competition for Synaptic Seeding Factors. Dev Cell 50, 447–461.e8. 10.1016/j.devcel.2019.06.014.

12. Kiral, F.R., Linneweber, G.A., Mathejczyk, T., Georgiev, S.V., Wernet, M.F., Hassan, B.A., von Kleist, M., and Hiesinger, P.R. (2020). Autophagy-dependent filopodial kinetics restrict synaptic partner choice during Drosophila brain wiring. Nat Commun 11, 1325. 10.1038/s41467-020-14781-4.

13. Lapraz, F., Boutres, C., Fixary-Schuster, C., De Queiroz, B.R., Plaçais, P.Y., Cerezo, D., Besse, F., Préat, T., and Noselli, S. (2023). Asymmetric activity of NetrinB controls laterality of the Drosophila brain. Nature Communications 2023 14:1 14, 1–13. 10.1038/S41467-023-36644-4.

14. GA, L., M, A., SB, D., M, B., L, H., G, L., RK, E., AD, S., M, W., PR, H., et al. (2020). A neurodevelopmental origin of behavioral individuality in the Drosophila visual system. Science 367, 1105–1112. 10.1126/SCIENCE.AAW7182.

15. Meng, J.L., Wang, Y., Carrillo, R.A., and Heckscher, E.S. (2020). Temporal transcription factors determine circuit membership by permanently altering motor neuron-to-muscle synaptic partnerships. Elife 9, 1–23. 10.7554/ELIFE.56898.

16. Klingler, E., Tomasello, U., Prados, J., Kebschull, J.M., Contestabile, A., Galiñanes, G.L., Fièvre, S., Santinha, A., Platt, R., Huber, D., et al. (2021). Temporal controls over inter-areal cortical projection neuron fate diversity. Nature 2021 599:7885 599, 453– 457. 10.1038/S41586-021-04048-3.

17. Jain, S., Lin, Y., Kurmangaliyev, Y.Z., Valdes-Aleman, J., LoCascio, S.A., Mirshahidi, P., Parrington, B., and Zipursky, S.L. (2022). A global timing mechanism regulates cell-type-specific wiring programmes. Nature 2022 603:7899 603, 112–118. 10.1038/S41586-022-04418-5.

18. Seroka, A.Q., and Doe, C.Q. (2019). The hunchback temporal transcription factor determines motor neuron axon and dendrite targeting in drosophila. Development (Cambridge) 146. 10.1242/DEV.175570/VIDEO-2.

19. Misgeld, T., and Schwarz, T.L. (2017). Mitostasis in Neurons: Maintaining Mitochondria in an Extended Cellular Architecture. Neuron 96, 651–666. 10.1016/J.NEURON.2017.09.055.

20. Kiral, F.R., Linneweber, G.A., Mathejczyk, T., Georgiev, S.V., Wernet, M.F., Hassan, B.A., von Kleist, M., and Hiesinger, P.R. (2020). Autophagy-dependent filopodial kinetics restrict synaptic partner choice during Drosophila brain wiring. Nature Communications 2020 11:1 11, 1–14. 10.1038/s41467-020-14781-4.

21. Menon, S., and Gupton, S.L. (2016). Building Blocks of Functioning Brain: Cytoskeletal Dynamics in Neuronal Development. Int Rev Cell Mol Biol 322, 183–245. 10.1016/BS.IRCMB.2015.10.002.

22. Rangaraju, V., Lewis, T.L., Hirabayashi, Y., Bergami, M., Motori, E., Cartoni, R., Kwon, S.K., and Courchet, J. (2019). Pleiotropic mitochondria: The influence of mitochondria on neuronal development and disease. Journal of Neuroscience 39, 8200–8208. 10.1523/JNEUROSCI.1157-19.2019.

23. Pekkurnaz, G., and Wang, X. (2022). Mitochondrial heterogeneity and homeostasis through the lens of a neuron. Nat Metab 4, 802–812. 10.1038/s42255-022-00594-w.

24. Verstreken, P., Ly, C. V., Venken, K.J.T., Koh, T.W., Zhou, Y., and Bellen, H.J. (2005). Synaptic mitochondria are critical for mobilization of reserve pool vesicles at Drosophila neuromuscular junctions. Neuron 47, 365–378. 10.1016/j.neuron.2005.06.018.

25. Lewis, T.L., Kwon, S.K., Lee, A., Shaw, R., and Polleux, F. (2018). MFF-dependent mitochondrial fission regulates presynaptic release and axon branching by limiting axonal mitochondria size. Nat Commun 9. 10.1038/s41467-018-07416-2.

26. Rangaraju, V., Lewis, T.L., Hirabayashi, Y., Bergami, M., Motori, E., Cartoni, R., Kwon, S.K., and Courchet, J. (2019). Pleiotropic Mitochondria: The Influence of Mitochondria on Neuronal Development and Disease. Journal of Neuroscience 39, 8200–8208. 10.1523/JNEUROSCI.1157-19.2019.

27. Faits, M.C., Zhang, C., Soto, F., and Kerschensteiner, D. (2016). Dendritic mitochondria reach stable positions during circuit development. Elife 5. 10.7554/ELIFE.11583.

28. Courchet, J., Lewis, T.L., Lee, S., Courchet, V., Liou, D.Y., Aizawa, S., and Polleux, F. (2013). Terminal axon branching is regulated by the LKB1-NUAK1 kinase pathway via presynaptic mitochondrial capture. Cell 153, 1510. 10.1016/J.CELL.2013.05.021.

29. Pickles, S., Vigié, P., and Youle, R.J. (2018). Mitophagy and Quality Control Mechanisms in Mitochondrial Maintenance. Current Biology 28, R170–R185. 10.1016/J.CUB.2018.01.004.

30. Cheng, X.T., Huang, N., and Sheng, Z.H. (2022). Programming axonal mitochondrial maintenance and bioenergetics in neurodegeneration and regeneration. Neuron 110, 1899–1923. 10.1016/J.NEURON.2022.03.015.

31. Iwata, R., Casimir, P., Erkol, E., Boubakar, L., Planque, M., Gallego López, I.M., Ditkowska, M., Gaspariunaite, V., Beckers, S., Remans, D., et al. (2023). Mitochondria metabolism sets the species-specific tempo of neuronal development. Science (1979) 379. 10.1126/SCIENCE.ABN4705/SUPPL_FILE/SCIENCE.ABN4705_MDAR_REPRODU CIBILITY_CHECKLIST.PDF.

32. Malin, J., and Desplan, C. (2021). Neural specification, targeting, and circuit formation during visual system assembly. Proc Natl Acad Sci U S A 118, e2101823118. 10.1073/PNAS.2101823118/ASSET/65A6A51E-98FC-4889-9257-B636CF57BB4F/ASSETS/IMAGES/LARGE/PNAS.2101823118FIG04.JPG.

33. Srahna, M., Leyssen, M., Ching, M.C., Fradkin, L.G., Noordermeer, J.N., and Hassan, B.A. (2006). A signaling network for patterning of neuronal connectivity in the Drosophila brain. PLoS Biol 4, 2076–2090. 10.1371/JOURNAL.PBIO.0040348.

34. Davie, K., Janssens, J., Koldere, D., De Waegeneer, M., Pech, U., Kreft, Ł., Aibar, S., Makhzami, S., Christiaens, V., Bravo González-Blas, C., et al. (2018). A Single-Cell Transcriptome Atlas of the Aging Drosophila Brain. Cell 174, 982–998.e20. 10.1016/j.cell.2018.05.057.

35. Nichterwitz, S., Chen, G., Aguila Benitez, J., Yilmaz, M., Storvall, H., Cao, M., Sandberg, R., Deng, Q., and Hedlund, E. (2016). ARTICLE Laser capture microscopy coupled with Smart-seq2 for precise spatial transcriptomic profiling. Nat Commun. 10.1038/ncomms12139.

36. Heo, Y.R., Lee, M.H., Kwon, S.Y., Cho, J., and Lee, J.H. (2019). TZAP mutation leads to poor prognosis of patients with breast cancer. Medicina (Lithuania) 55. 10.3390/medicina55110748.

37. Arantes dos Santos, G., Viana, N.I., Pimenta, R., Reis, S.T., Ramos Moreira Leite, K., and Srougi, M. (2021). Hypothesis: The triad androgen receptor, zinc finger proteins and telomeres modulates the global gene expression pattern during prostate cancer progression. Med Hypotheses 150. 10.1016/J.MEHY.2021.110566.

38. Özel, M.N., Simon, F., Jafari, S., Holguera, I., Chen, Y.C., Benhra, N., El-Danaf, R.N., Kapuralin, K., Malin, J.A., Konstantinides, N., et al. (2020). Neuronal diversity and convergence in a visual system developmental atlas. Nature 2020 589:7840 589, 88– 95. 10.1038/S41586-020-2879-3.

39. Port, F., Strein, C., Stricker, M., Rauscher, B., Heigwer, F., Zhou, J., Beyersdörffer, C., Frei, J., Hess, A., Kern, K., et al. A large-scale resource for tissue-specific CRISPR mutagenesis in Drosophila. 10.1101/636076.

40. Su, J., Li, Z., Fusté, J.M., Simavorian, T., Bartocci, C., Tsai, J., Karlseder, J., and Denchi, E.L. TZAP: A telomere-associated protein involved in telomere length control.

41. Kudron, M.M., Victorsen, A., Gevirtzman, L., Hillier, L.W., Fisher, W.W., Vafeados, D., Kirkey, M., Hammonds, A.S., Gersch, J., Ammouri, H., et al. (2018). The ModERN Resource: Genome-Wide Binding Profiles for Hundreds of Drosophila and Caenorhabditis elegans Transcription Factors. Genetics 208, 937–949. 10.1534/GENETICS.117.300657.

42. Jahn, A., Rane, G., Paszkowski-Rogacz, M., Sayols, S., Bluhm, A., Han, C., Draškovič, I., Londoño-Vallejo, J.A., Kumar, A.P., Buchholz, F., et al. (2017). ZBTB 48 is both a vertebrate telomere-binding protein and a transcriptional activator. EMBO Rep 18, 929–946. 10.15252/embr.201744095.

43. Detmer, S.A., and Chan, D.C. (2007). Functions and dysfunctions of mitochondrial dynamics. Nat Rev Mol Cell Biol 8, 870–879. 10.1038/nrm2275.

44. Ahmad, T., Aggarwal, K., Pattnaik, B., Mukherjee, S., Sethi, T., Tiwari, B.K., Kumar, M., Micheal, A., Mabalirajan, U., Ghosh, B., et al. (2013). Computational classification of mitochondrial shapes reflects stress and redox state. Cell Death Dis 4. 10.1038/cddis.2012.213.

45. Chan, D.C. (2006). Mitochondria: Dynamic Organelles in Disease, Aging, and Development. Cell 125, 1241–1252. 10.1016/j.cell.2006.06.010.

46. Gomes, L.C., Benedetto, G. Di, and Scorrano, L. (2011). During autophagy mitochondria elongate, are spared from degradation and sustain cell viability. Nat Cell Biol 13, 589–598. 10.1038/ncb2220.

47. Ashrafi, G., de Juan-Sanz, J., Farrell, R.J., and Ryan, T.A. (2020). Molecular Tuning of the Axonal Mitochondrial Ca2+ Uniporter Ensures Metabolic Flexibility of Neurotransmission. Neuron 105, 678–687.e5. 10.1016/J.NEURON.2019.11.020.

48. Cuhadar, U., Calzado-Reyes, L., Pascual-Caro, C., Aberra, A.S., Aggarwal, A., Podgorski, K., Hoppa, M.B., and Juan-Sanz, J. de (2022). Activity-driven synaptic translocation of LGI1 controls excitatory neurotransmission. bioRxiv, 2022.07.03.498586. 10.1101/2022.07.03.498586.

49. Jin, E.J., Kiral, F.R., Ozel, M.N., Burchardt, L.S., Osterland, M., Epstein, D., Wolfenberg, H., Prohaska, S., and Hiesinger, P.R. (2018). Live Observation of Two Parallel Membrane Degradation Pathways at Axon Terminals. Curr Biol 28, 1027–1038.e4. 10.1016/J.CUB.2018.02.032.

50. Talay, M., Richman, E.B., Snell, N.J., Hartmann, G.G., Fisher, J.D., Sorkaç, A., Santoyo, J.F., Chou-Freed, C., Nair, N., Johnson, M., et al. (2017). Transsynaptic Mapping of Second-Order Taste Neurons in Flies by trans-Tango. Neuron 96, 783–795.e4. 10.1016/j.neuron.2017.10.011.

51. Dutta, S., Linneweber, G.A., Andriatsilavo, M., Hiesinger, P.R., and Hassan, B.A. (2022). A Critical Developmental Interval of Coupling Axon Branching to Synaptic Degradation During Neural Circuit Formation. SSRN Electronic Journal. 10.2139/SSRN.4076344.

52. Sweeney, S.T., Broadie, K., Keane, J., Niemann, H., and O’Kane, C.J. (1995). Targeted expression of tetanus toxin light chain in Drosophila specifically eliminates synaptic transmission and causes behavioral defects. Neuron 14, 341–351. 10.1016/0896-6273(95)90290-2.

53. Hodge, J.J.L. (2009). Ion channels to inactivate neurons in Drosophila. Front Mol Neurosci 2. 10.3389/NEURO.02.013.2009.

54. Nitabach, M.N., Blau, J., and Holmes, T.C. (2002). Electrical silencing of Drosophila pacemaker neurons stops the free-running circadian clock. Cell 109, 485–495. 10.1016/S0092-8674(02)00737-7.

55. Colomb, J., Reiter, L., Blaszkiewicz, J., Wessnitzer, J., and Brembs, B. (2012). Open source tracking and analysis of adult Drosophila locomotion in Buridan’s paradigm with and without visual targets. PLoS One 7. 10.1371/JOURNAL.PONE.0042247.

56. Bonnefoy, E., Orsi, G.A., Couble, P., and Loppin, B. (2007). The Essential Role of Drosophila HIRA for De Novo Assembly of Paternal Chromatin at Fertilization. PLoS Genet 3, 1991–2006. 10.1371/JOURNAL.PGEN.0030182.

57. Loppin, B., Bonnefoy, E., Anselme, C., Laurençon, A., Karr, T.L., and Couble, P. (2005). The histone H3.3 chaperone HIRA is essential for chromatin assembly in the male pronucleus. Nature 437, 1386–1390. 10.1038/NATURE04059.

58. Ray-Gallet, D., Quivy, J.P., Scamps, C., Martini, E.M.D., Lipinski, M., and Almouzni, G. (2002). HIRA is critical for a nucleosome assembly pathway independent of DNA synthesis. Mol Cell 9, 1091–1100. 10.1016/S1097-2765(02)00526-9.

59. Rust, K., Tiwari, M.D., Mishra, V.K., Grawe, F., and Wodarz, A. (2018). Myc and the Tip60 chromatin remodeling complex control neuroblast maintenance and polarity in Drosophila. EMBO J 37, e98659. 10.15252/EMBJ.201798659.

60. Cenci, G., Ciapponi, L., and Gatti, M. (2005). The mechanism of telomere protection: A comparison between Drosophila and humans. Chromosoma 114, 135–145. 10.1007/s00412-005-0005-9.

61. Louis, E.J. (2002). Are Drosophila telomeres an exception or the rule?

62. Groza, T., Lopez Gomez, F., Mashhadi, H.H., Muñoz, V., Muñoz-Fuentes, M., Gunes, O., Wilson, R., Cacheiro, P., Frost, A., Keskivali-Bond, P., et al. (2023). The International Mouse Phenotyping Consortium: comprehensive knockout phenotyping underpinning the study of human disease. Nucleic Acids Res 51, D1038–D1045. 10.1093/NAR/GKAC972.

63. Fahrner, J.A., Liu, R., Perry, M.S., Klein, J., and Chan, D.C. (2016). A novel de novo dominant negative mutation in DNM1L impairs mitochondrial fission and presents as childhood epileptic encephalopathy. Am J Med Genet A 170, 2002–2011. 10.1002/ajmg.a.37721.

64. Romero-Garcia, S., and Prado-Garcia, H. (2019). Mitochondrial calcium: Transport and modulation of cellular processes in homeostasis and cancer (Review). Int J Oncol 54, 1155–1167. 10.3892/IJO.2019.4696/HTML.

65. White, R.J., and Reynolds, I.J. (1997). Mitochondria accumulate Ca2+ following intense glutamate stimulation of cultured rat forebrain neurones. J Physiol 498, 31. 10.1113/JPHYSIOL.1997.SP021839.

66. Redpath, C.J., Bou Khalil, M., Drozdzal, G., Radisic, M., and McBride, H.M. (2013). Mitochondrial hyperfusion during oxidative stress is coupled to a dysregulation in calcium handling within a C2C12 cell model. PLoS One 8. 10.1371/JOURNAL.PONE.0069165.

67. Deas, E., Plun-Faureau, H., and Wood, N.W. (2009). PINK1 function in health and disease. EMBO Mol Med 1, 152. 10.1002/EMMM.200900024.

68. Heeman, B., Van den Haute, C., Aelvoet, S.A., Valsecchi, F., Rodenburg, R.J., Reumers, V., Debyser, Z., Callewaert, G., Koopman, W.J.H., Willems, P.H.G.M., et al. (2011). Depletion of PINK1 affects mitochondrial metabolism, calcium homeostasis and energy maintenance. J Cell Sci 124, 1115–1125. 10.1242/JCS.078303.

69. Gao, X., Yu, X., Zhang, C., Wang, Y., Sun, Y., Sun, H., Zhang, H., Shi, Y., and He, X. (2022). Telomeres and Mitochondrial Metabolism: Implications for Cellular Senescence and Age-related Diseases. Stem Cell Rev Rep 18, 2315–2327. 10.1007/S12015-022-10370-8.

70. Nassour, J., Aguiar, L.G., Correia, A., Schmidt, T.T., Mainz, L., Przetocka, S., Haggblom, C., Tadepalle, N., Williams, A., Shokhirev, M.N., et al. (2023). Telomere-to-mitochondria signalling by ZBP1 mediates replicative crisis. Nature 2023, 1–7. 10.1038/s41586-023-05710-8.

71. Iyer, E.P.R., and Cox, D.N. (2010). Laser capture Microdissection of Drosophila peripheral neurons. Journal of Visualized Experiments. 10.3791/2016.

72. Dobin, A., Davis, C.A., Schlesinger, F., Drenkow, J., Zaleski, C., Jha, S., Batut, P., Chaisson, M., and Gingeras, T.R. (2013). STAR: ultrafast universal RNA-seq aligner. Bioinformatics 29, 15–21. 10.1093/bioinformatics/bts635.

73. Plaçais, P.Y., De Tredern, É., Scheunemann, L., Trannoy, S., Goguel, V., Han, K.A., Isabel, G., and Preat, T. (2017). Upregulated energy metabolism in the Drosophila mushroom body is the trigger for long-term memory. Nature Communications 2017 8:1 8, 1–14. 10.1038/ncomms15510.

74. R Core Team (2022). R: A Language and Environment for Statistical Computing.

75. Legros, F., Lombès, A., Frachon, P., and Rojo, M. (2002). Mitochondrial Fusion in Human Cells Is Efficient, Requires the Inner Membrane Potential, and Is Mediated by Mitofusins. Mol Biol Cell 13, 4343–4354. 10.1091/mbc.e02-06-0330.

76. Sankaranarayanan, S., De Angelis, D., Rothman, J.E., and Ryan, T.A. (2000). The Use of pHluorins for Optical Measurements of Presynaptic Activity. Biophys J 79, 2199–2208. 10.1016/S0006-3495(00)76468-X.

77. Aggarwal, A., Liu, R., Chen, Y., Ralowicz, A.J., Bergerson, S.J., Tomaska, F., Mohar, B., Hanson, T.L., Hasseman, J.P., Reep, D., et al. (2023). Glutamate indicators with improved activation kinetics and localization for imaging synaptic transmission. Nat Methods. 10.1038/s41592-023-01863-6.

78. Merrill, R.A., Flippo, K.H., and Strack, S. (2017). Measuring Mitochondrial Shape with ImageJ. In, pp. 31–48. 10.1007/978-1-4939-6890-9_2.

